# Analysis and Exploration of the Implementation System of Mixed Ownership of Scientific and Technological Achievements in Colleges and Universities

**DOI:** 10.1101/2022.09.04.506550

**Authors:** Panjun Gao, Yong Qi, Qing Guo

## Abstract

Due to the backwardness of the management model and the rigidity of the system, the current service invention system restricts the transformation of scientific and technological achievements of colleges and universities to a certain extent. The mixed ownership system of university post achievements, which gives universities and inventors the freedom to dispose of the rights of scientific and technological achievements, and fundamentally stimulates the enthusiasm of all the achievement owners to transform their achievements, is an important exploration to promote the transformation of scientific and technological achievements. However, there are still many problems in the implementation of the mixed ownership of scientific and technological achievements in colleges and universities. Through the questionnaire survey and analysis, it is found that the four problems such as defects in the management system for the distribution of rights and interests of scientific and technological achievements, imperfect evaluation indicators of scientific and technological achievements, imbalance in the service system for the transformation of scientific and technological achievements, and lack of authoritative standards for the evaluation of the value of the transformation of scientific and technological achievements are the main problems hindering the development of mixed ownership of the current college post achievements. Based on the existing problems, it is proposed to continue to improve the management system, assessment system and service system of mixed ownership of scientific and technological achievements in colleges and universities. The research results can provide new evidence and policy enlightenment for the development of mixed ownership of scientific and technological achievements in colleges and universities in China.

## Introduction

With the increasing intensity of competition between countries, the scientific and technological achievements of universities have become an important force in improving the country’s comprehensive competitive strength. The developed countries have not only generally established a relatively complete system of laws and regulations on the transfer and transformation of scientific and technological achievements, but also formulated the corresponding implementation rules to promote the transfer and transformation of various scientific and technological achievements. The common law system represented by the United States and the civil law system represented by Germany and Japan have made detailed and clear provisions on the transfer and transformation of scientific and technological achievements in colleges and universities, property rights recognition rules, and the distribution of benefits, etc. Under the legal framework of various countries, they have not lost the unity of ‘flexibility’ and ‘timeliness’. which worth us to take its essence from its dross. According to China’s 2020 patent survey and statistics, the industrialization rate of effective invention patents in China’s colleges and universities in 2020 is 3.8% [1], and the industrialization rate of invention patents in China’s colleges and universities is lower than that of the survey data released in Japan (patent utilization rate of 47.6% [2]). Due to the particularity of China’s socialist mechanism, the introduction of the system of scientific and technological achievements in western countries is not conducive to the transfer and transformation of scientific and technological achievements in China’s colleges and universities. Therefore, at present, all colleges and universities in China are actively exploring the transformation mode of achievements adapted to the environment of China’s mechanism. With the optimization of the transformation system of scientific and technological achievements by relevant departments of the state, the mixed ownership of post scientific and technological achievements has become a research hotspot in the field of domestic scientific and technological achievements transformation.

There is still some controversy about whether the mixed ownership of scientific and technological achievements can promote the transfer and transformation of scientific and technological achievements. Some scholars believe that the mixed ownership of post scientific and technological achievements does not conform to the relevant legislative provisions of China’s current scientific and technological achievements, and whether it can mobilize the enthusiasm of inventors has not been fully confirmed. If the interest distribution incentive mechanism of the mixed ownership of post scientific and technological achievements is unreasonable, it may hinder the transformation of scientific and technological achievements. However, most scholars have positive opinions on the mixed ownership of post scientific and technological achievements, and generally believe that the mixed ownership of scientific and technological achievements in colleges and universities determines the property rights of scientific and technological achievements from the source, ensures the interests of all aspects of the main body of scientific and technological achievements, and is conducive to improving the transformation momentum of scientific and technological achievements of colleges and universities.

### Research on the mixed ownership of post scientific and technological achievements

Some domestic colleges and universities have implemented the pilot reform of mixed ownership of post science and technology achievements, and also proposed implementation models and mechanisms that can promote the transformation of scientific and technological achievements. At the same time, relevant domestic laws and regulations are also actively adjusting to promote the reasonable compliance of mixed ownership of post scientific and technological achievements. Tang Liangzhi (2014) believes that in accordance with the principle of “who completes, who owns, and who benefits”, the property rights relationship between universities and scientific researchers should be reasonably defined, a system for the transfer and transformation of scientific and technological achievements with unified responsibilities and rights should be established, and a pilot project of transferring the ownership of scientific and technological achievements to the inventors of scientific and technological achievements [3]. Kang Kaining (2015) believes that mixed ownership is an effective means to promote the transformation of scientific and technological achievements, which solves the problems of market-oriented pricing of post scientific and technological achievements, equity awards when evaluating the price of shares, and the preservation and appreciation of the state-owned assets in post scientific and technological achievements, and the mixed ownership system makes the scientific research direction and scientific research topic selection change to market-oriented [4]. Liu Feng et al. (2017) analyzed the mixed ownership system of post scientific and technological achievements from the perspective of property rights theory, and believes that giving inventors and units the right to agree on the ownership of achievements would help reduce transaction costs in the process of transformation of achievements and enhance the motivation of inventors to engage in the transformation of achievements [5]. Zhang Mingshen (2017) found that by recognizing the special attributes of post scientific and technological achievements and implementing the incentive of ownership confirmation, etc., the “state-owned asset curse” can be broken, and the incentives of scientific researchers and universities in power and motivation can be realized [6]. Pan Xiaoyu et al. (2017) found that after Southwest Jiaotong University innovatively proposed the mixed ownership of post scientific and technological achievements, it also introduced implementation rules to analyze potential problems such as school control, school authorship, school intervention, and operation convenience in the process of transforming post scientific and technological achievements, which is conducive to promoting the transformation of scientific and technological achievements [7]. Yao Yang (2017) believes that the mixed production of scientific and technological achievements in colleges and universities is a new thing bred by reform practice, on the one hand, it makes rigid public property rights into clear mixed property rights; on the other hand, the effective participation of inventors of service scientific and technological achievements can improve the probability of successful transformation of scientific and technological achievements, and the practical effect is significant [8]. Zhang Wenfei (2019) conducts research on the new ownership model of mixed ownership of post scientific and technological achievements from the perspective of the change of tenure system based on post scientific and technological achievements by using economic analysis methods and the investment costs of employees and employers as an analysis tool. He makes the optimal system selection according to the results of the “cost-benefit” game, and finds that the application of mixed ownership of post scientific and technological achievements can produce the expected benefits of reducing transaction costs, avoiding commons tragedies, and preventing the loss of state-owned assets [9]. Su Jicheng et al. (2019) believe that the current management model of post scientific and technological achievements is rigid and the legal system is lagging behind, which greatly limits the enthusiasm of scientific researchers to innovate and also restricts the transformation of scientific and technological achievements. The implementation of the “mixed ownership” reform of post scientific and technological achievements can solve the problem of idle and wasteful scientific and technological achievements of universities and scientific research institutes by giving play to the incentive function of the property rights market, and promote the institutional innovation of the transformation of the results market [10]. Gao Yanqiong et al. (2021) believe that the mixed ownership of scientific and technological achievements in colleges and universities can break down the existing institutional obstacles and fundamentally stimulate the transformation enthusiasm of all the people who achieve results, which is of great promotion significance [11].

### Research on the mixed ownership of post scientific and technological achievements

Although some regions have put forward relevant rules, policies, and implementation rules in the process of implementing the pilot reform of mixed ownership of scientific and technological achievements, they are still not very suitable for the current laws and regulations in China from the overall level, and whether the mixed ownership of job scientific and technological achievements can promote the transformation of scientific and technological achievements has not really been proved by practice. For example, Chen Baiqiang et al. (2017) believe that the mixed ownership of job scientific and technological achievements violates laws and regulations such as the Patent Law, is not necessary from the actual need to promote the transformation of scientific and technological achievements, and is extremely unreasonable, and may also lead to the diversification and complexity of the use and disposal of post scientific and technological achievements at the operational level, hindering the transformation and implementation of scientific and technological achievements [12]. Wu Shouren (2018) believes that the mixed ownership of post scientific and technological achievements is illegal, unreasonable, inconvenient to operate and unnecessary, and suggests a simplistic treatment, that is, post scientific and technological achievements should belong to the unit, scientific and technological personnel cannot enjoy the ownership of job scientific and technological achievements, including applying for intellectual property rights with job scientific and technological achievements, and the person who completes the post scientific and technological achievements cannot be registered as the intellectual property rights holder [13].

### Research on the neutral view of mixed ownership of post scientific and technological achievements

Most scholars maintain a neutral attitude towards the mixed ownership of post scientific and technological achievements, and believe that the implementation of mixed ownership of scientific and technological achievements has certain practical significance, but it is still necessary to improve and make breakthroughs in models and mechanisms. For example, Xia Shuang (2018) believes that the lack of impartiality that may arise in the implementation of mixed ownership and controversy, such as: whether the distribution ratio is fair and reasonable, whether the policy should continue to encourage after reaching phased results, and how to protect state-owned assets behind the emphasis on the fairness of the rights and interests of subjects. It also used the relevant knowledge such as the theory of justice and the theory of responsibility ethics to analyze, think and summarize this, and put forward suggestions on the introduction of ethical regulations, system improvement, security system construction, government macro management and other aspects, and improved the mixed ownership of post scientific and technological achievements, so that they can play a full role and promote the rapid and smooth transformation of scientific and technological achievements [14]. Yao Yang (2018) believes that laws and policies show the characteristics of deregulation, decentralization, and the transformation and expansion of scientific and technological personnel, which form a favorable environment for the reform pilot of mixed ownership of post scientific and technological achievements, but the pilot reform also has the reality of confusing legal basis, and the continuous promotion of reform requires the authorization of the power organs or a clearer legal interpretation [15]. Li Tingting et al. (2020) believe that although the policy pilot of Sichuan Province has changed the property rights structure of universities in the province, it has not affected the increment, quality and transformation of patents significantly [16]. Wang Yinghang (2020) believes that the mixed ownership of scientific and technological achievements in colleges and universities is an important exploration for China to give scientific researchers the ownership of scientific and technological achievements and promote the transformation of scientific and technological achievements, but the current plan not only violates the provisions of the higher-level law on the ownership of post scientific and technological achievements, but also lacks a response plan to the inherent defects of mixed property rights [17].

In conclusion, the research on the mixed ownership of china’s post scientific and technological achievements, starting from 2015, with the pilot of the mixed ownership of post scientific and technological achievements in colleges and universities in the southwest region, many scholars hold positive opinions, but with the continuous implementation of the mixed ownership of post scientific and technological achievements in colleges and universities, it has not reached the anticipated goal. Therefore, since 2017, some scholars have begun to oppose it, believing that it is not necessary to implement the mixed ownership of post scientific and technological achievements in college and universities. Since 2018, with the in-depth analysis of the mixed ownership of scientific and technological achievements of university posts and put forward targeted countermeasures and suggestions, it is believed that after the improvement of the mixed ownership of scientific and technological achievements of college jobs, it is possible to achieve the expected goals. In the past two years, with the continuous improvement of the mixed ownership system of scientific and technological achievements in colleges and universities, some scholars have begun to hold affirmative opinions again.

There are many obstacles in promoting the mixed ownership of post scientific and technological achievements in colleges and universities, and it is still necessary to further improve the model and mechanism of the system. Currently, most scholars believe that the key to the implementation of the mixed ownership system of post scientific and technological achievements is the need to build a clear and reasonable ownership division and equity distribution mechanism, so that universities and inventors can maximize their interests and achieve the preservation and appreciation of state-owned assets in the true sense [18–21]. Therefore, in the context of the strategy of intellectual property power, it is urgent to conduct systematic and in-depth researches and analyses on the division of ownership and benefits of mixed ownership of scientific and technological achievements in colleges and universities in China, deepen the reform and innovation of the mixed ownership management mechanism for scientific and technological achievements of colleges and universities, use the cooperative game theory model to clarify the rationality of the implementation of the mixed ownership of scientific and technological achievements of colleges and universities, and promote the sustained and sound development of the mixed ownership of post scientific and technological achievements, which has key academic value and practical application prospects for promoting the transformation of scientific and technological achievements in China.

## Method

### Participant

China’s colleges and universities scientific and technological achievements have difficult to transform, low conversion rate, transformation of the benefits of creation is not prominent and other issues. To deal with these problems, some of China’s universities have introduced a pilot policy of mixed ownership of scientific and technological achievements, although certain results have been achieved, but still encountered obstacles in practice. Strengthening the study of mixed ownership of scientific and technological achievements in colleges and universities, especially related issues in the context of the system of post scientific and technological achievements, is conducive to promoting the transformation of scientific and technological achievements, and achieving the structural reform of the supply side of scientific and technological achievements.

After conducting in-depth discussions on the difficulties and pain points that hinder the ownership of scientific and technological achievements in colleges and universities and the reform of the benefit-sharing system, a questionnaire survey report was formulated. The main subjects of the survey is the scientific researchers and management personnel of the university, and a total of 340 valid questionnaires are collected.The basic information of participants is shown in Table 1. This study was approved by the Academic Committee of Nanjing University of Science and Technology, and informed written consent was obtained from all participants prior to actual participation. The survey questions and results are shown in figs 1-20.

**Fig 1.**
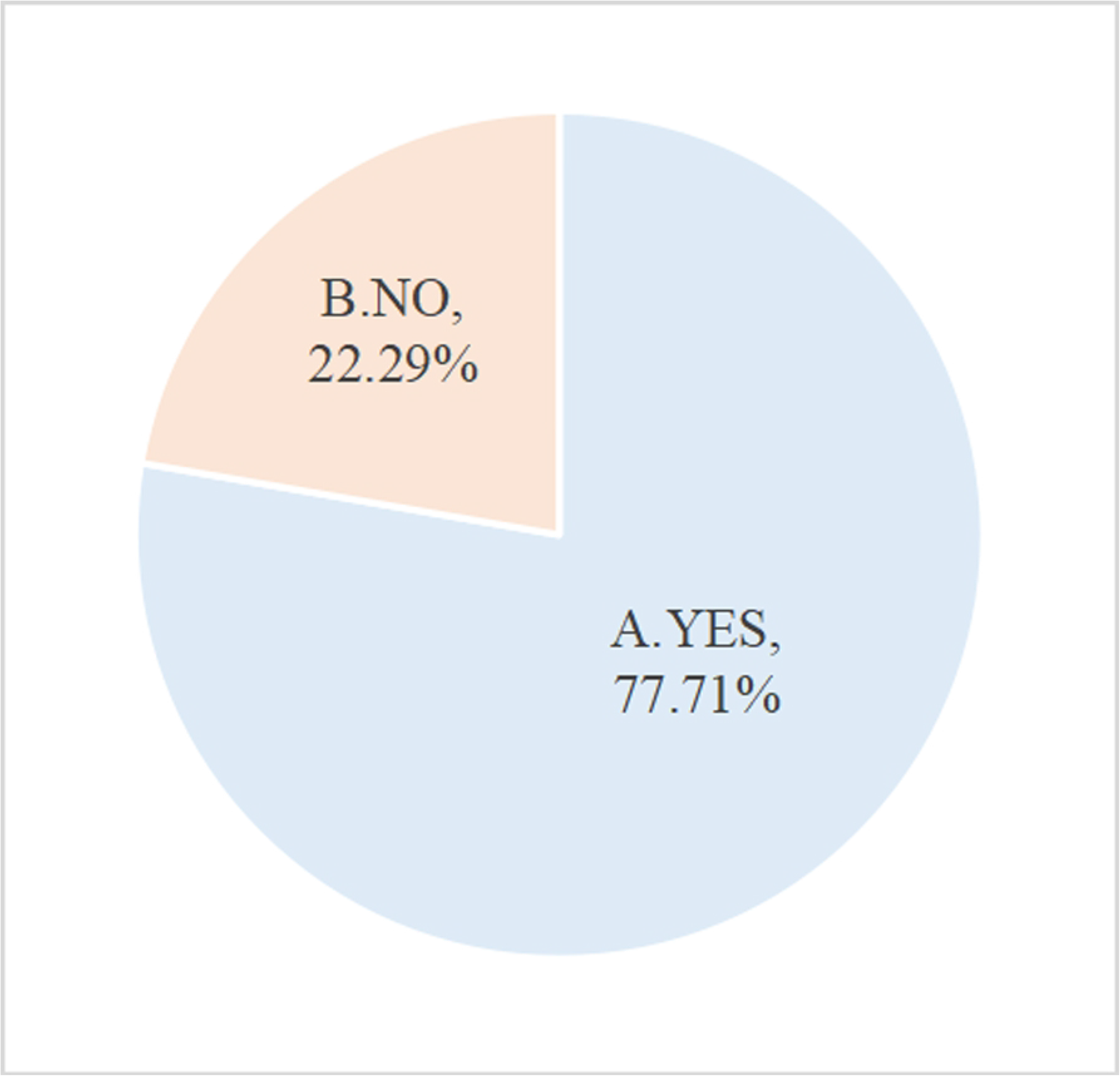
Has your university issued a managerial system for the distribution of RISTA? RISTA= rights and interests in science and technology achievements

**Fig 2.**
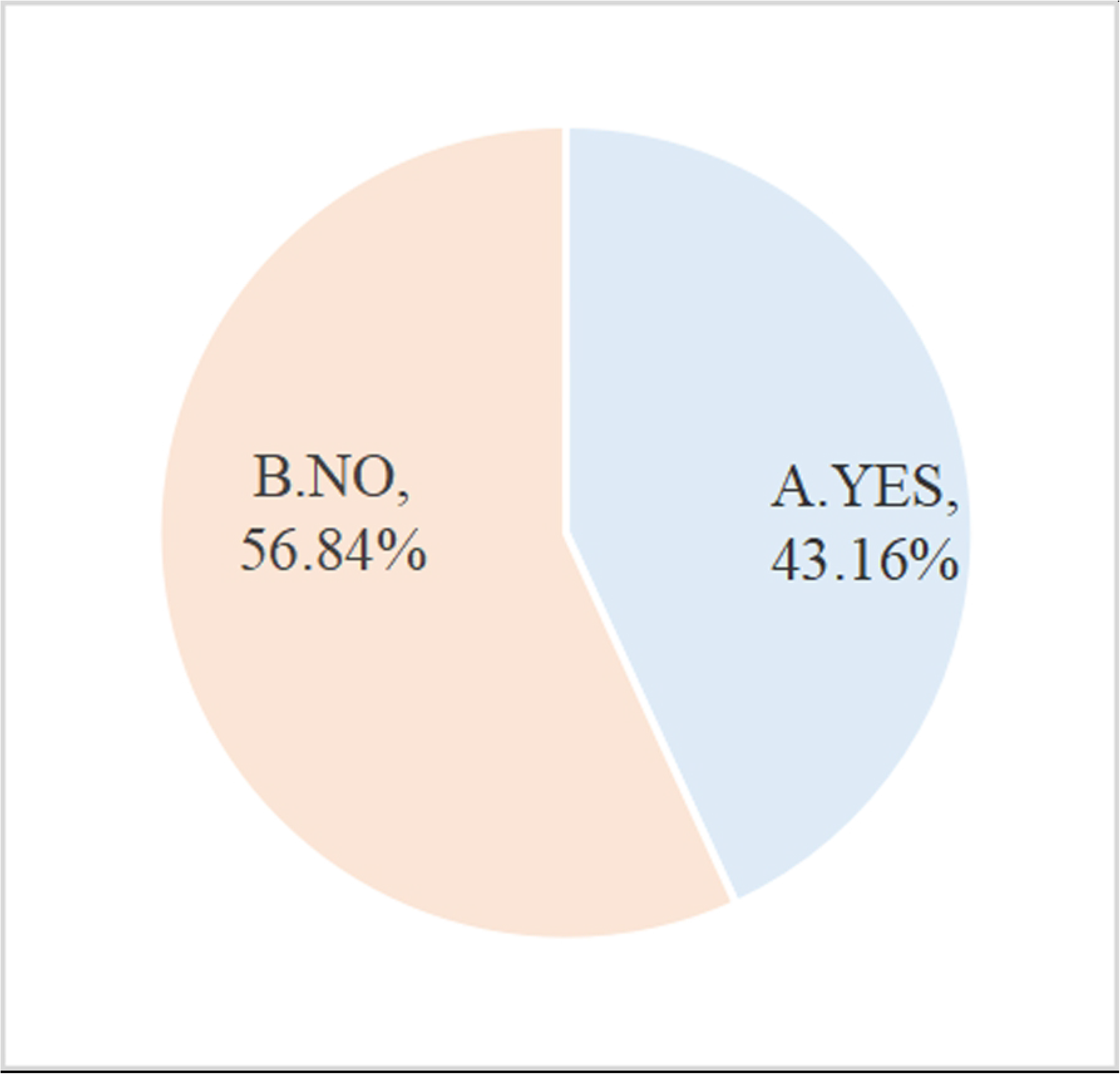
Do you know the managerial system for the RISTA issued by your university? RISTA= rights and interests in science and technology achievements

**Fig 3.**
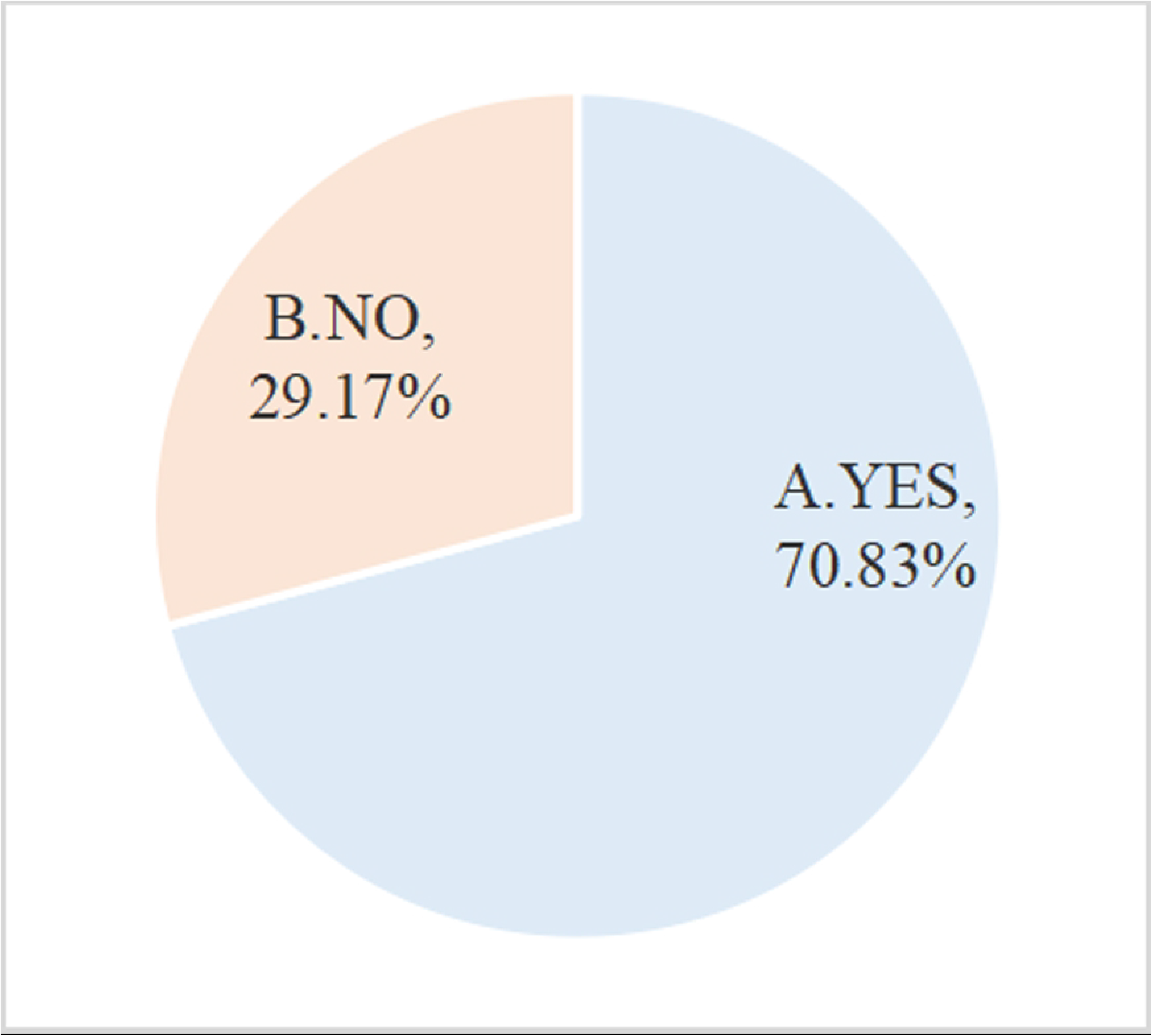
Has your university established a technology transfer agency or commissioned a third-party service agency?

**Fig 4.**
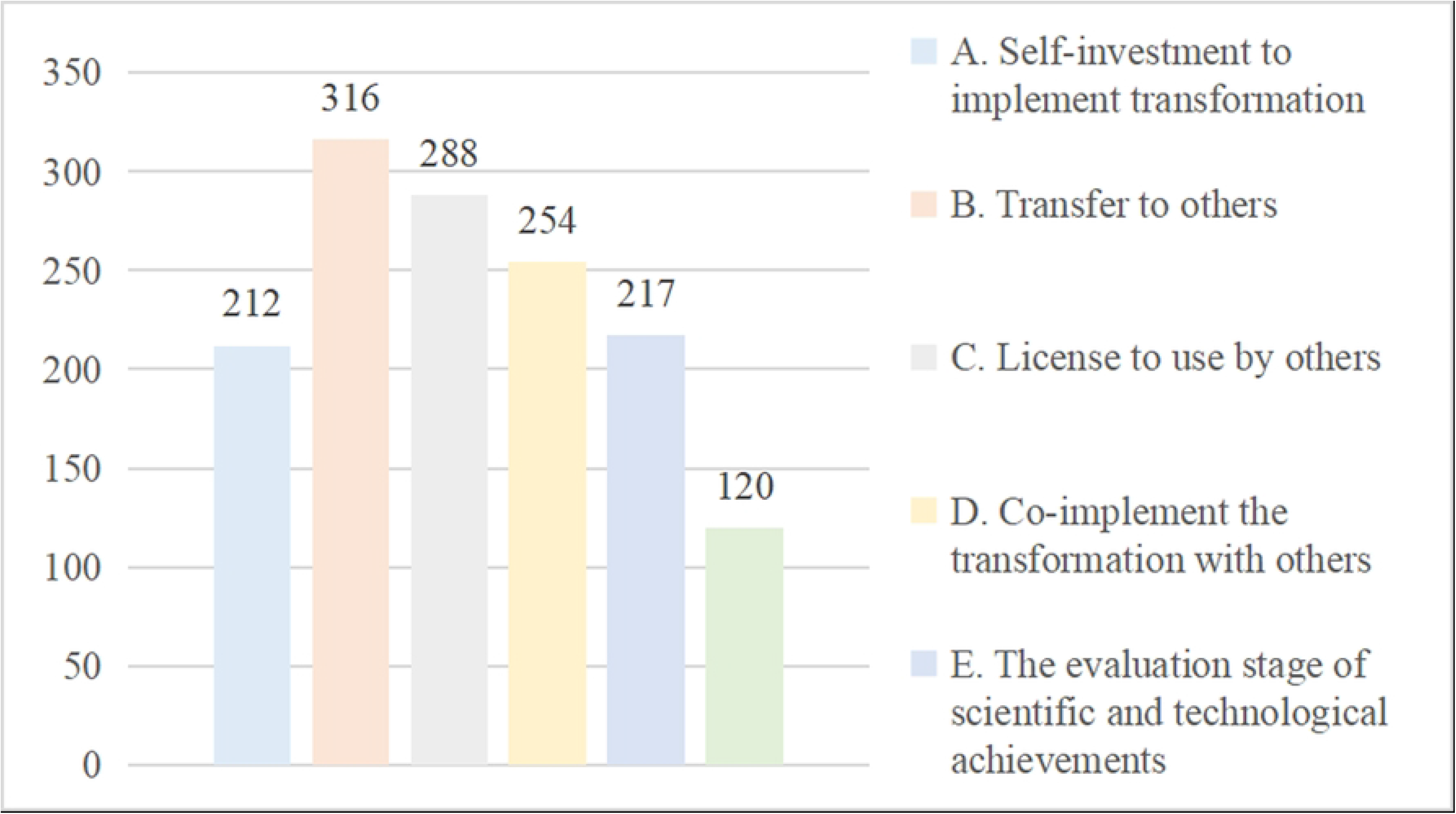
Which method does your university mainly use to transform STA? STA= science and technology achievements

**Fig 5.**
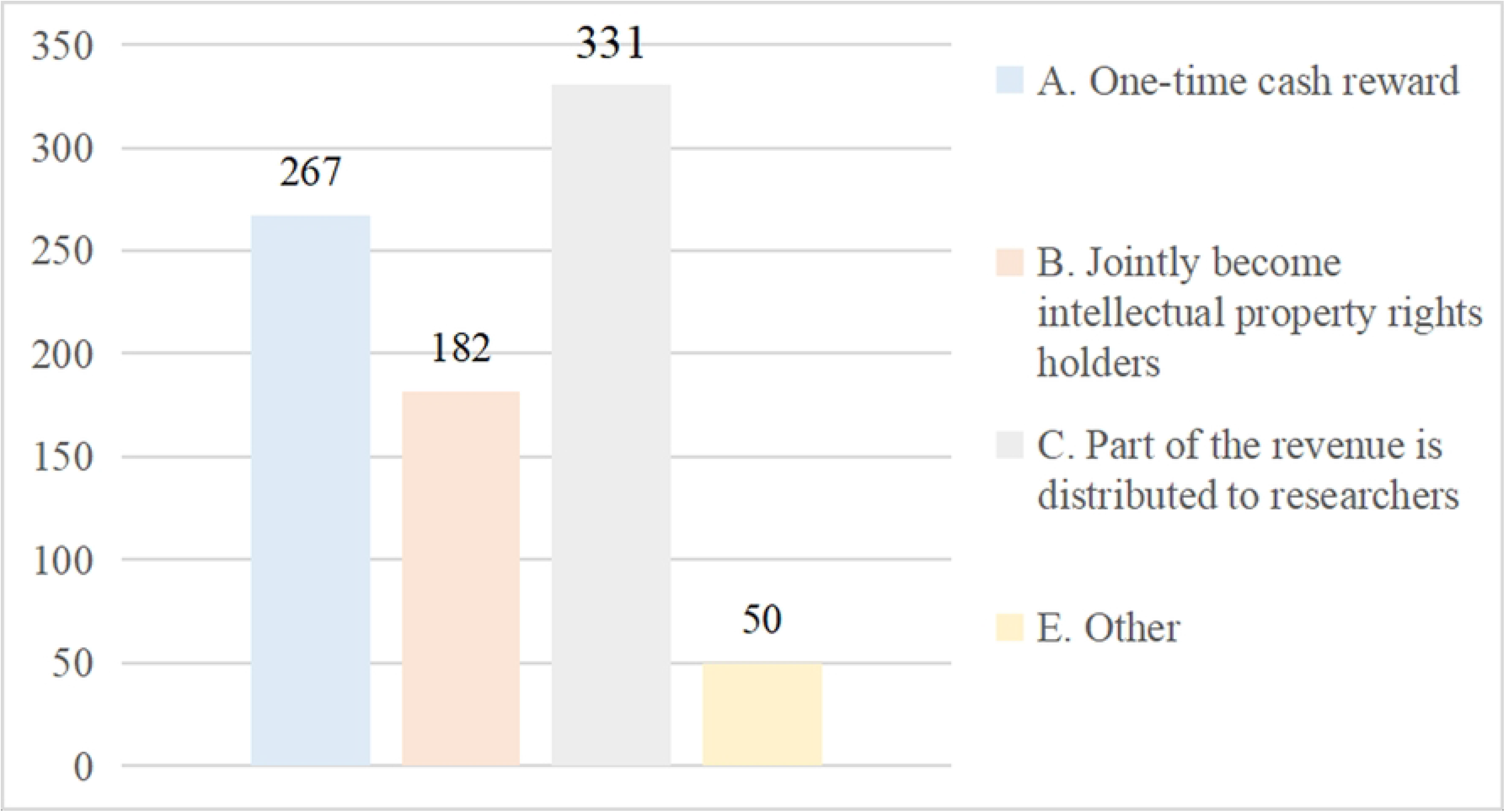
Which method of distribution of RISTA is mainly adopted by your university? RISTA=rights and interests in science and technology achievements

**Fig 6.**
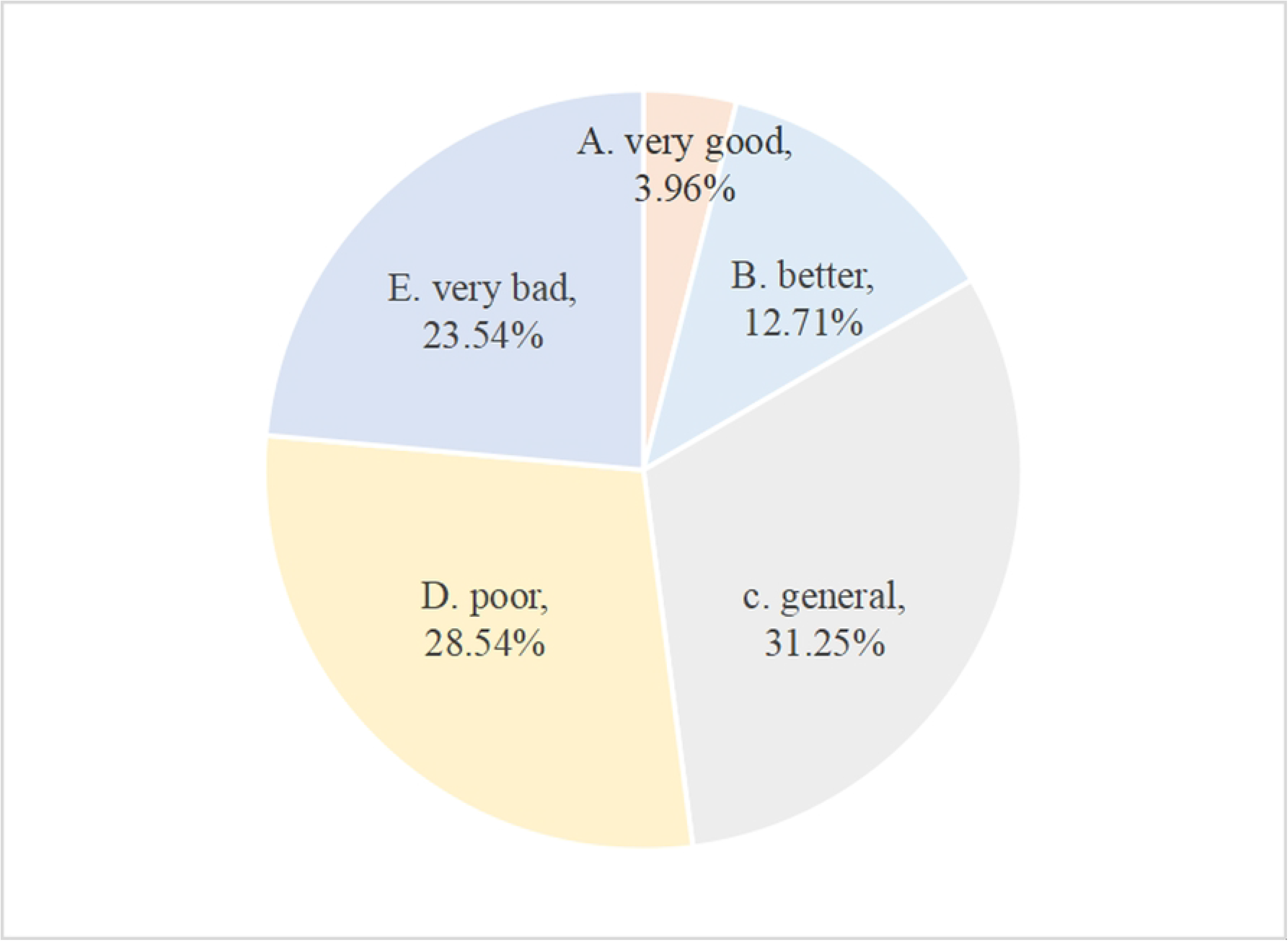
What is the incentive effect of your university’s measures for the distribution of RISTA ? RISTA=rights and interests in science and technology achievements

**Fig 7.**
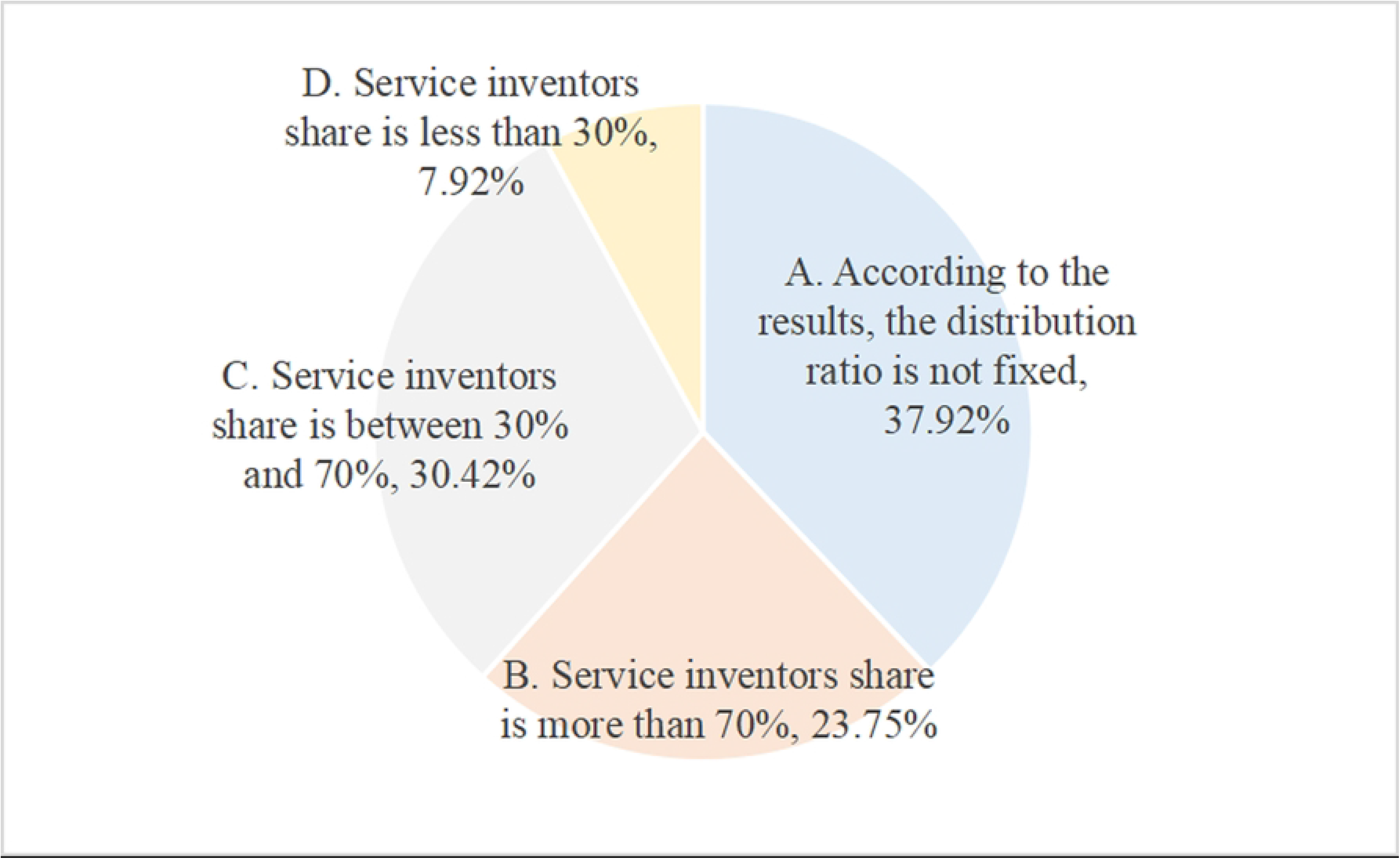
In terms of the distribution of RISTA, which distribution ratio does your university adopt? RISTA=rights and interests in science and technology achievements

**Fig 8.**
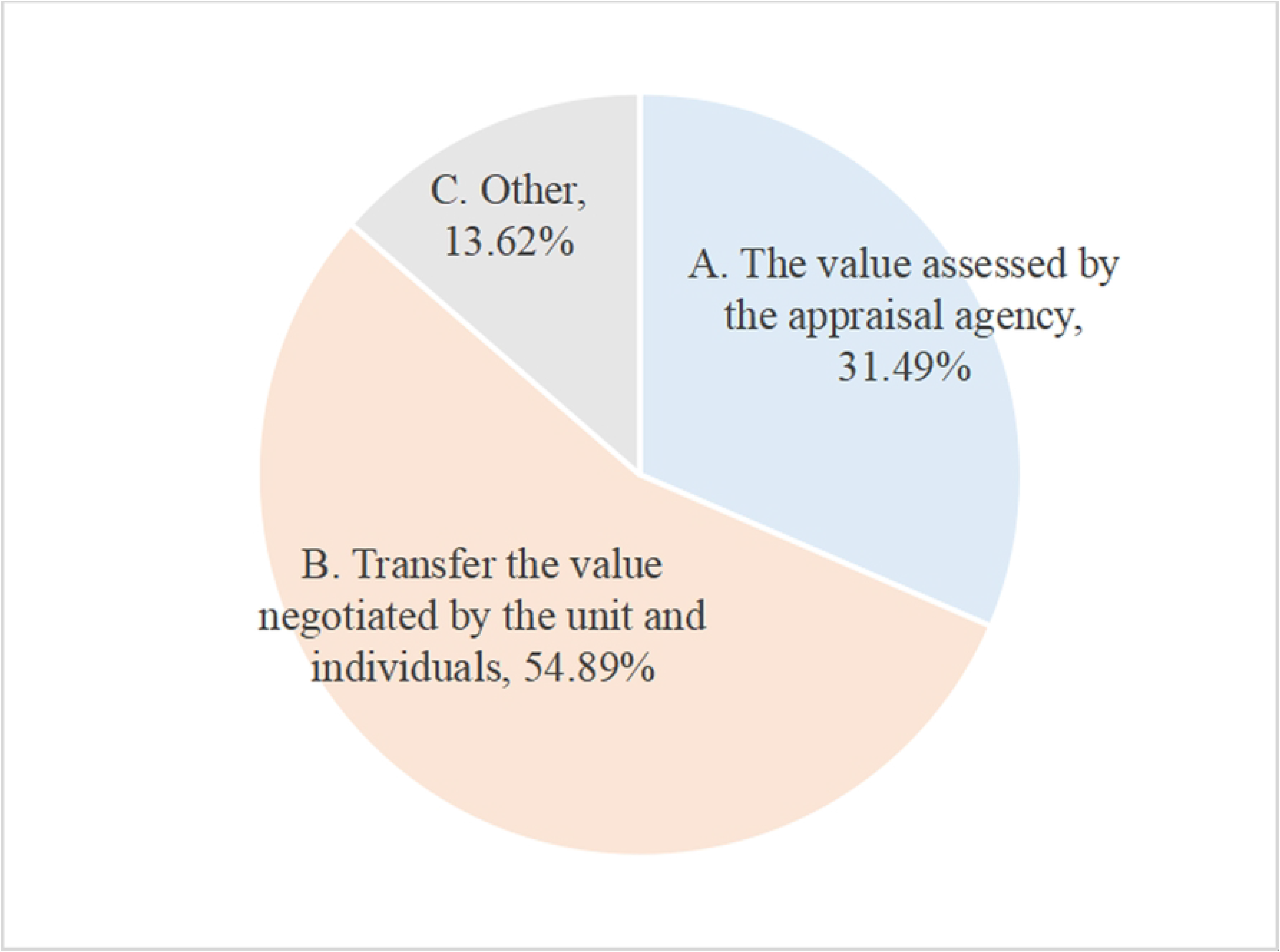
What is the most important way to determine the value of the transformation of STA? STA=science and technology achievements

**Fig 9.**
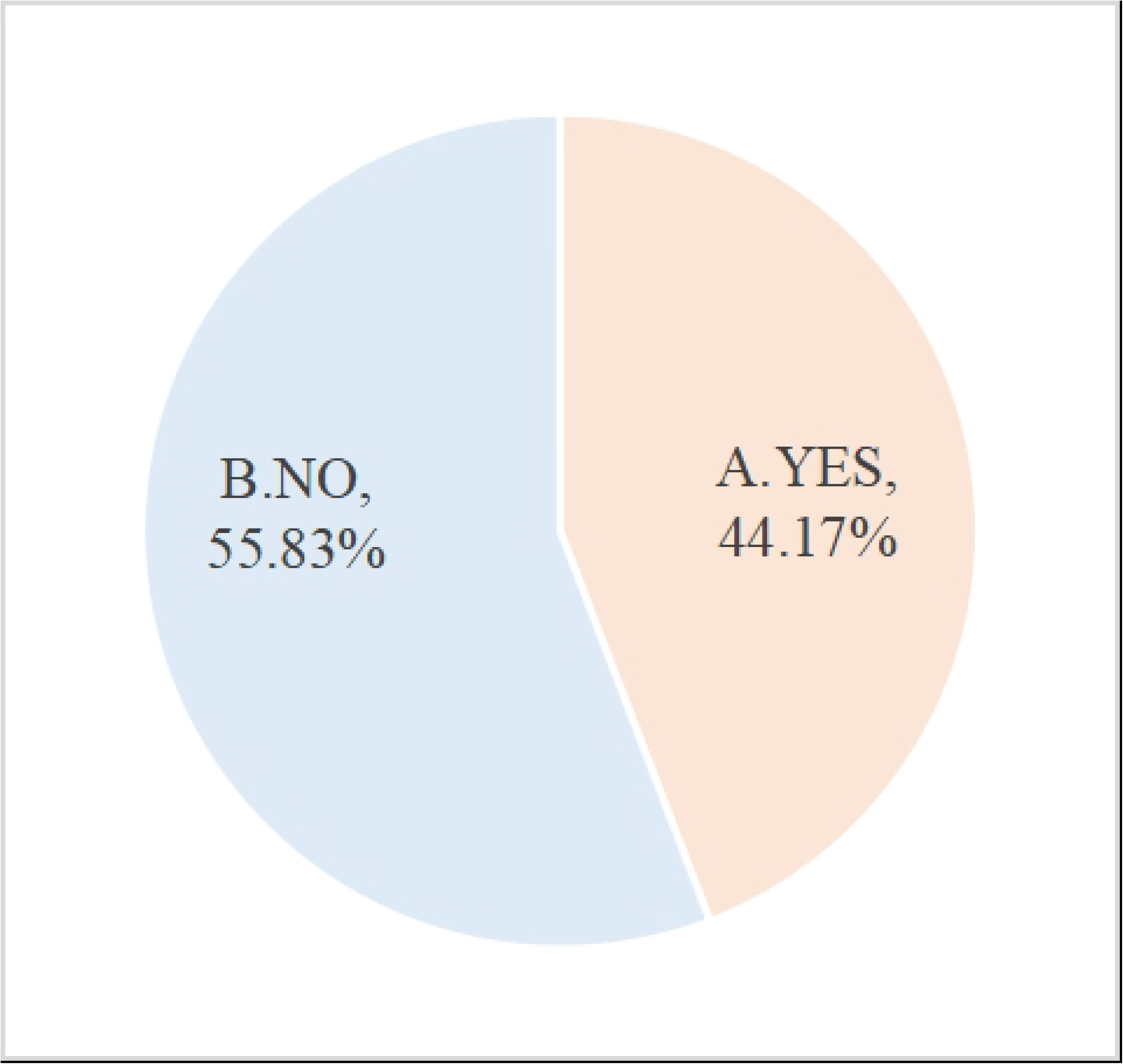
Does your university have STA value assessment institutions or entrust third-party institutions? STA=science and technology achievement

**Fig 10.**
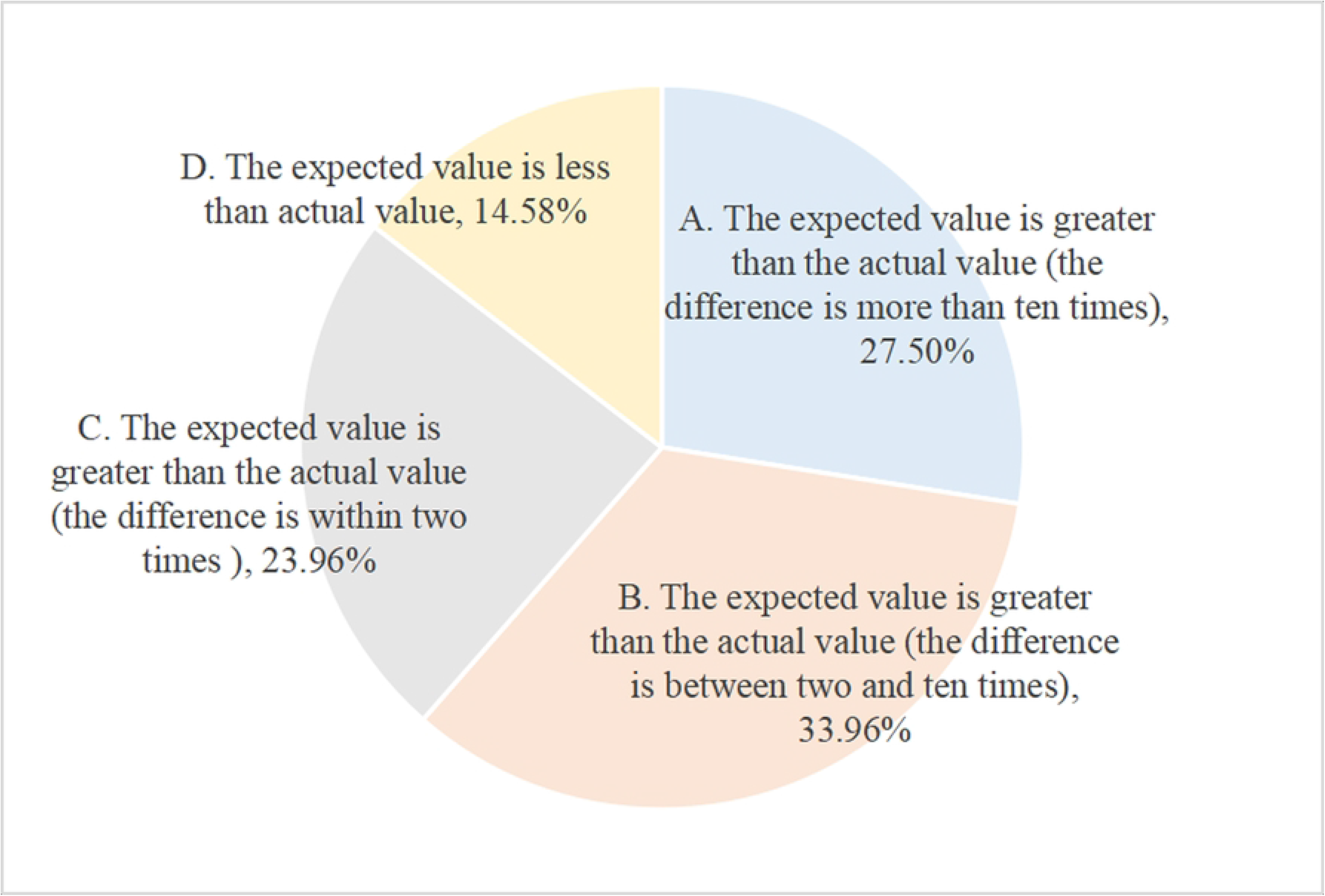
What is the relationship between expected value and actual value of STA ? STA= science and technology achievement

**Fig 11.**
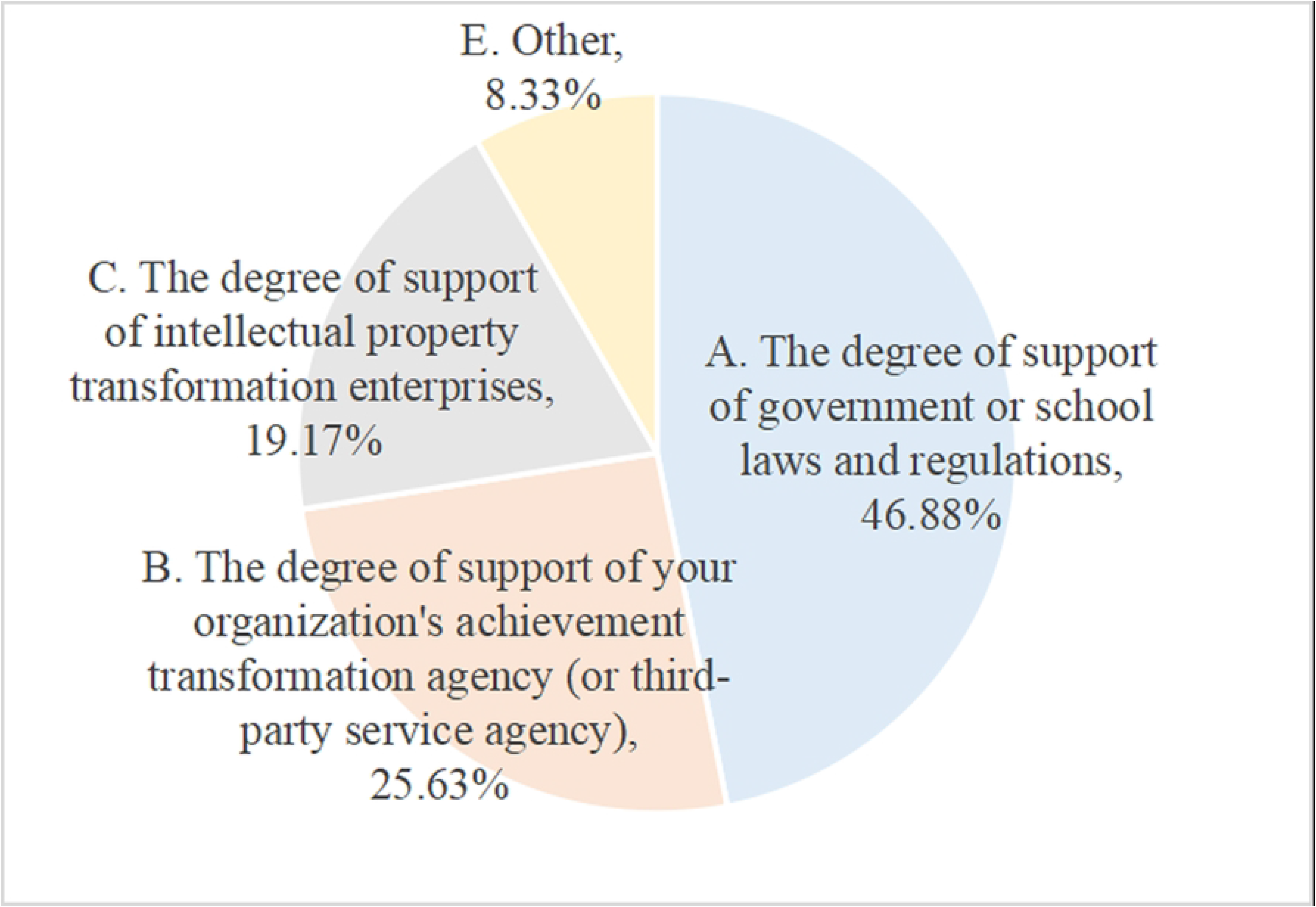
What is the reasons for the differences between expected value and actual value of STA ? STA=science and technology achievement

**Fig 12.**
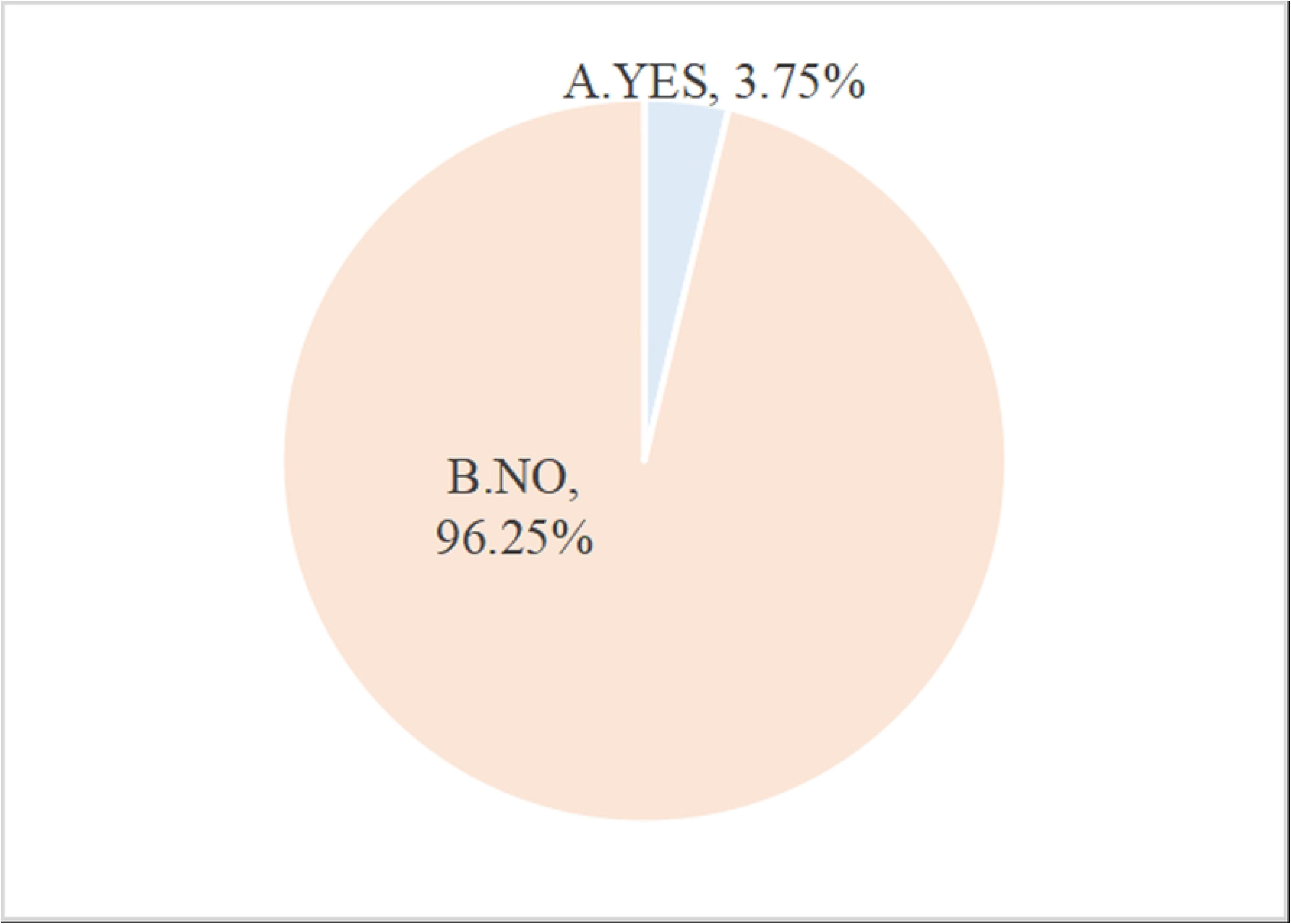
Has your university approved that the inventor of STA can be the right holder together with it? STA=science and technology achievement

**Fig 13.**
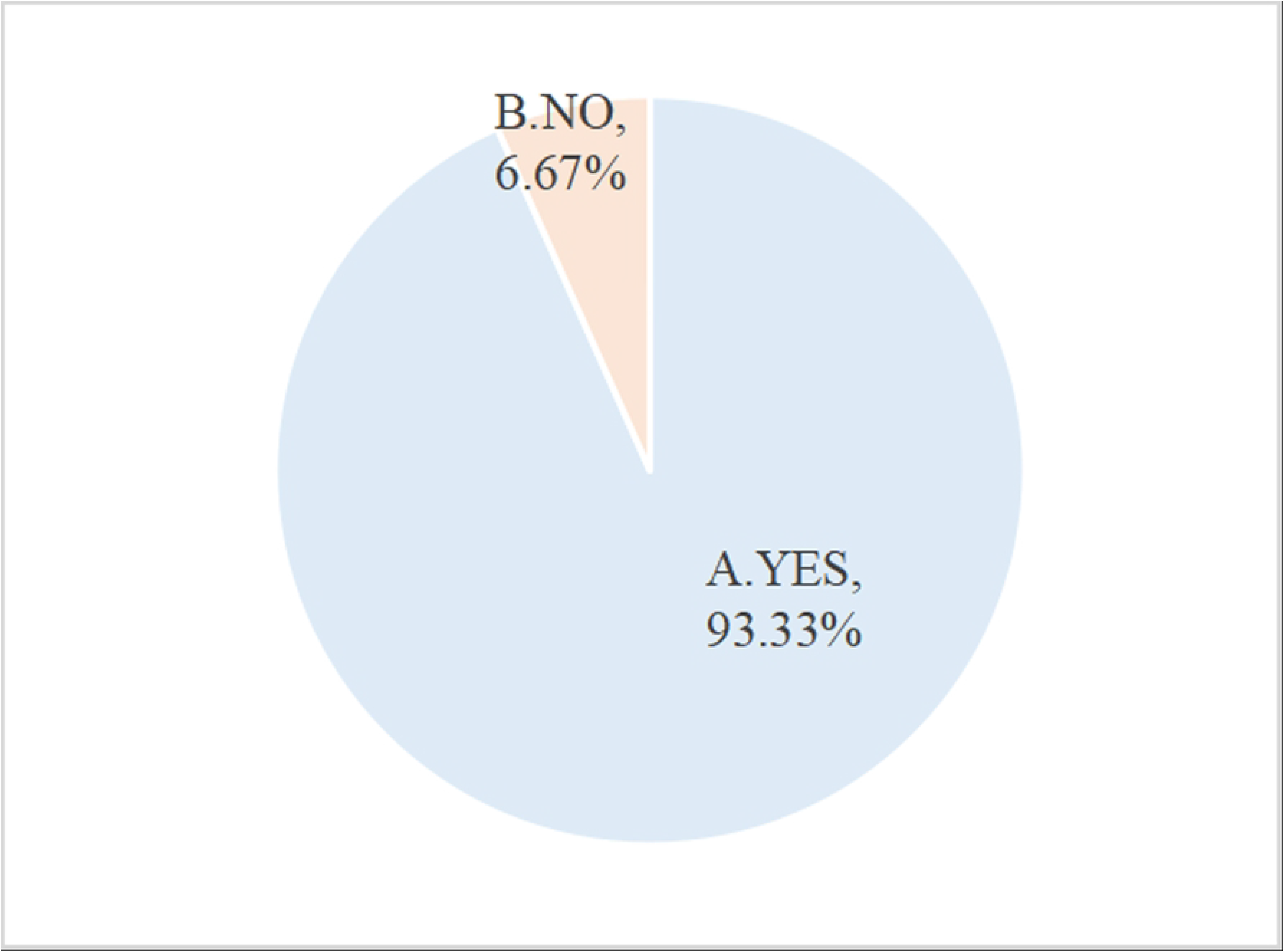
Is it necessary for inventors and university to jointly become right holders of STA? STA=science and technology achievement

**Fig 14.**
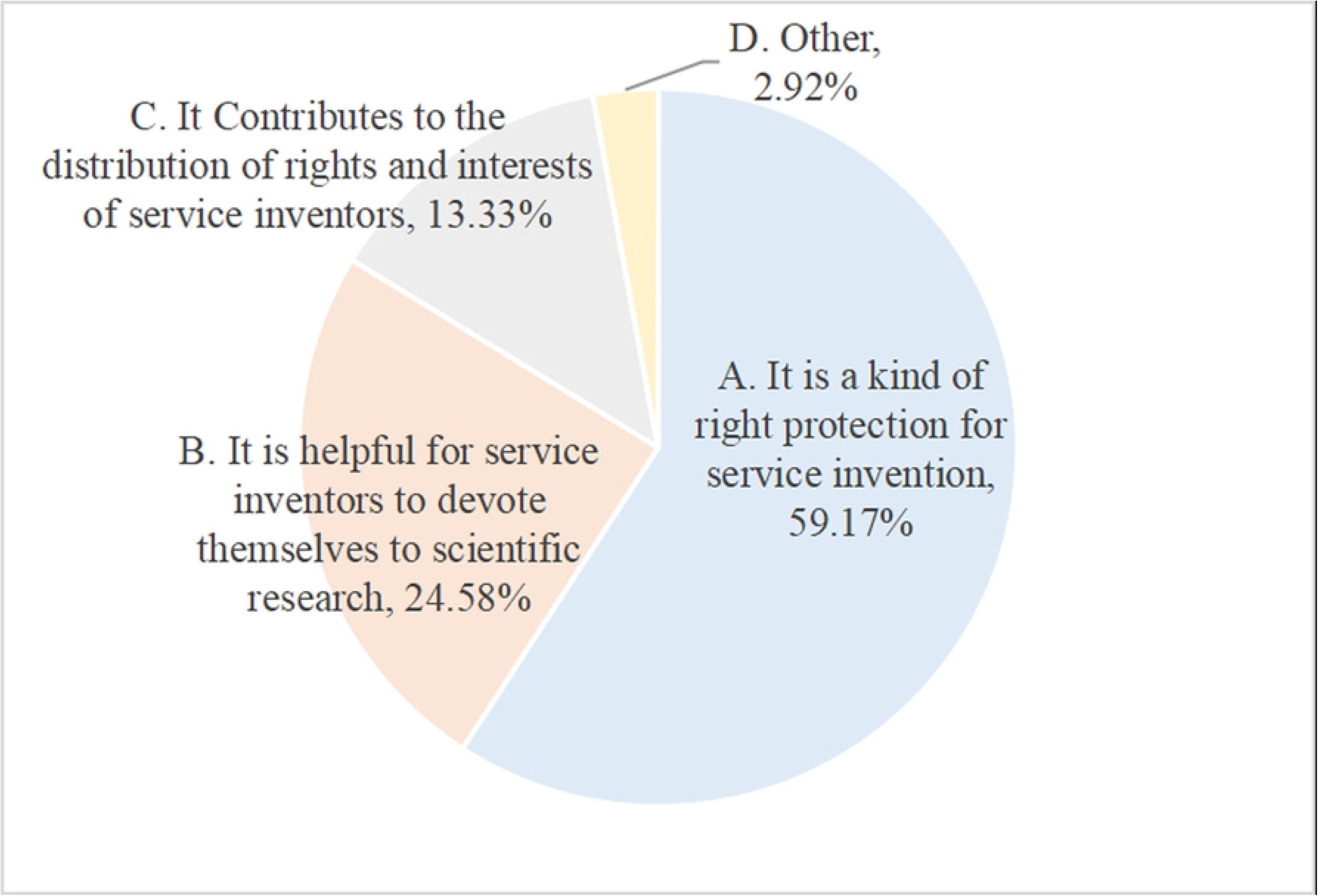
What is the reason for inventors and university to jointly become right holders of STA? STA=science and technology achievement

**Fig 15.**
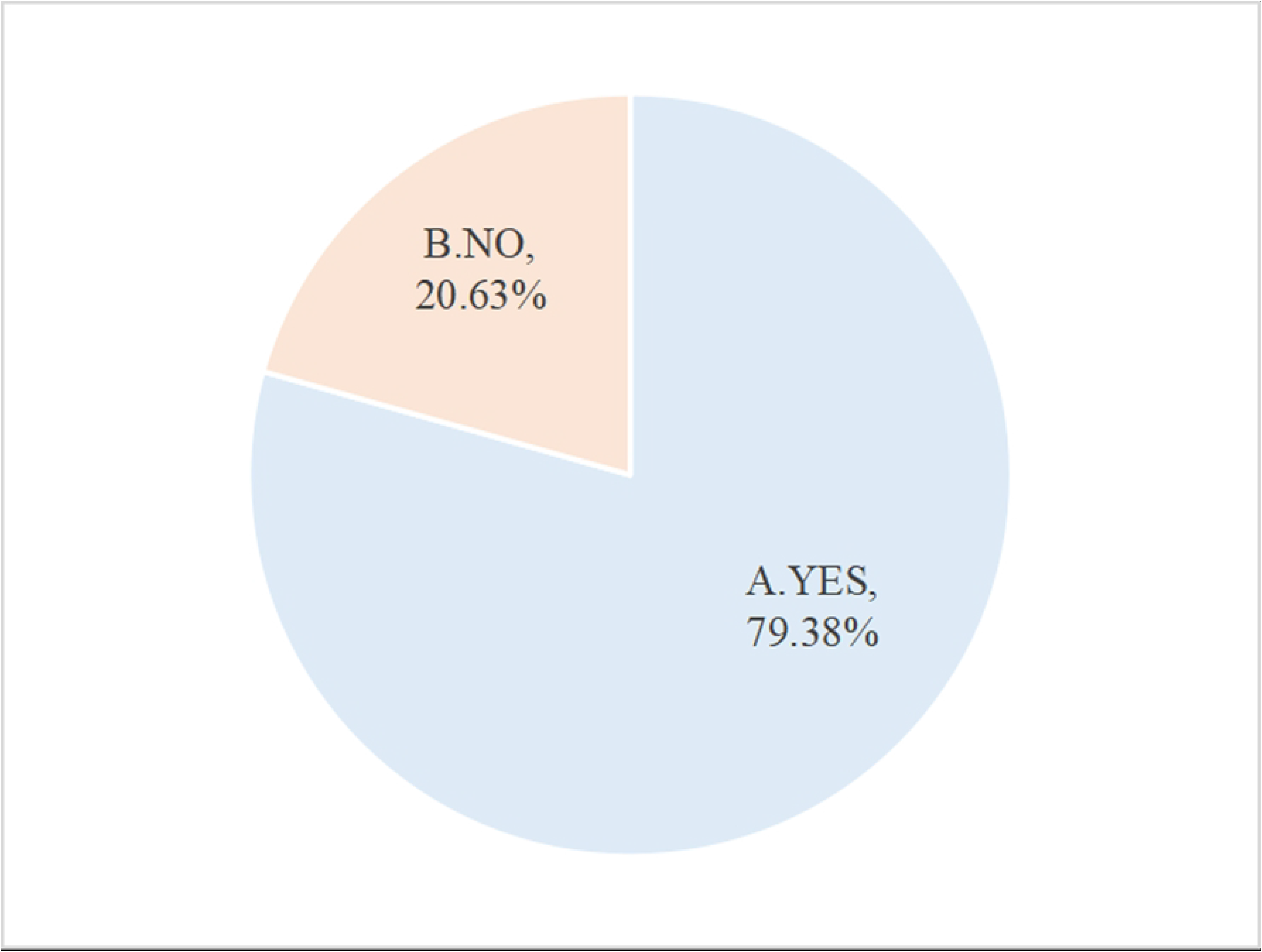
Are you willing to link the transformation effect of your STA with the assessment indicators? STA=science and technology achievement

**Fig 16.**
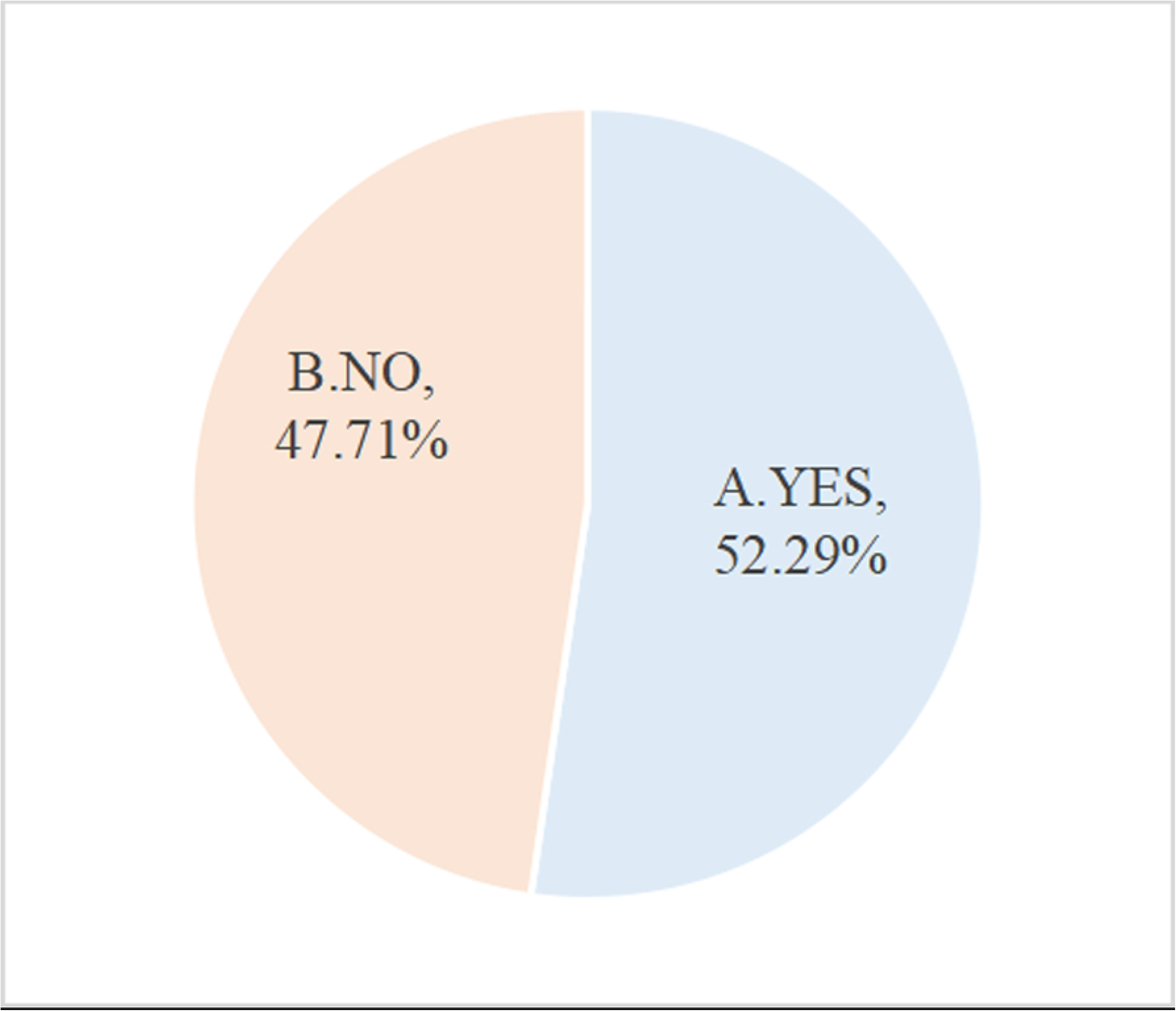
Does your university establish a transformation agency of STA or entrust a third-party agency? STA=science and technology achievement

**Fig 17.**
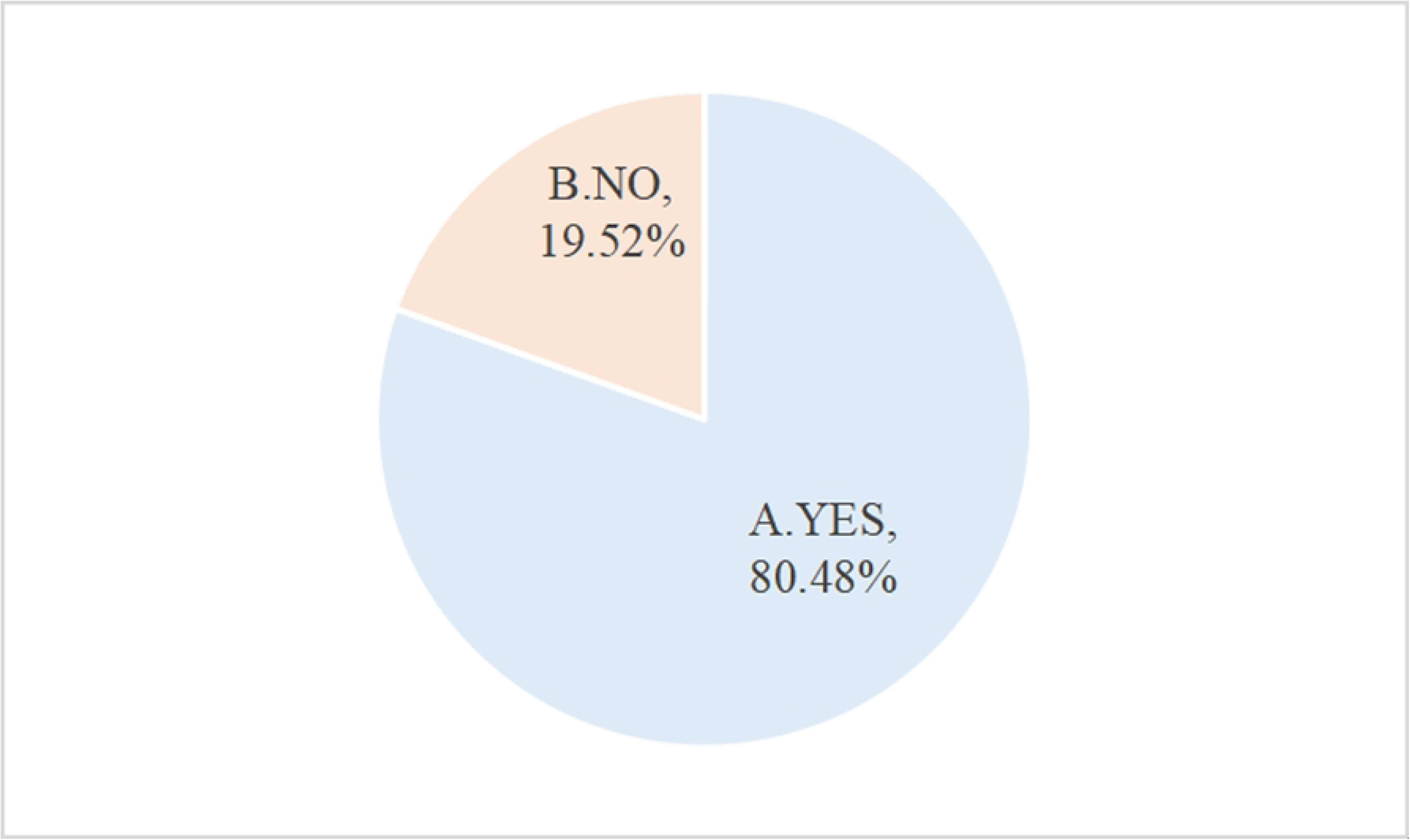
Are you satisfied with the achievement transformation service work of your university?

**Fig 18.**
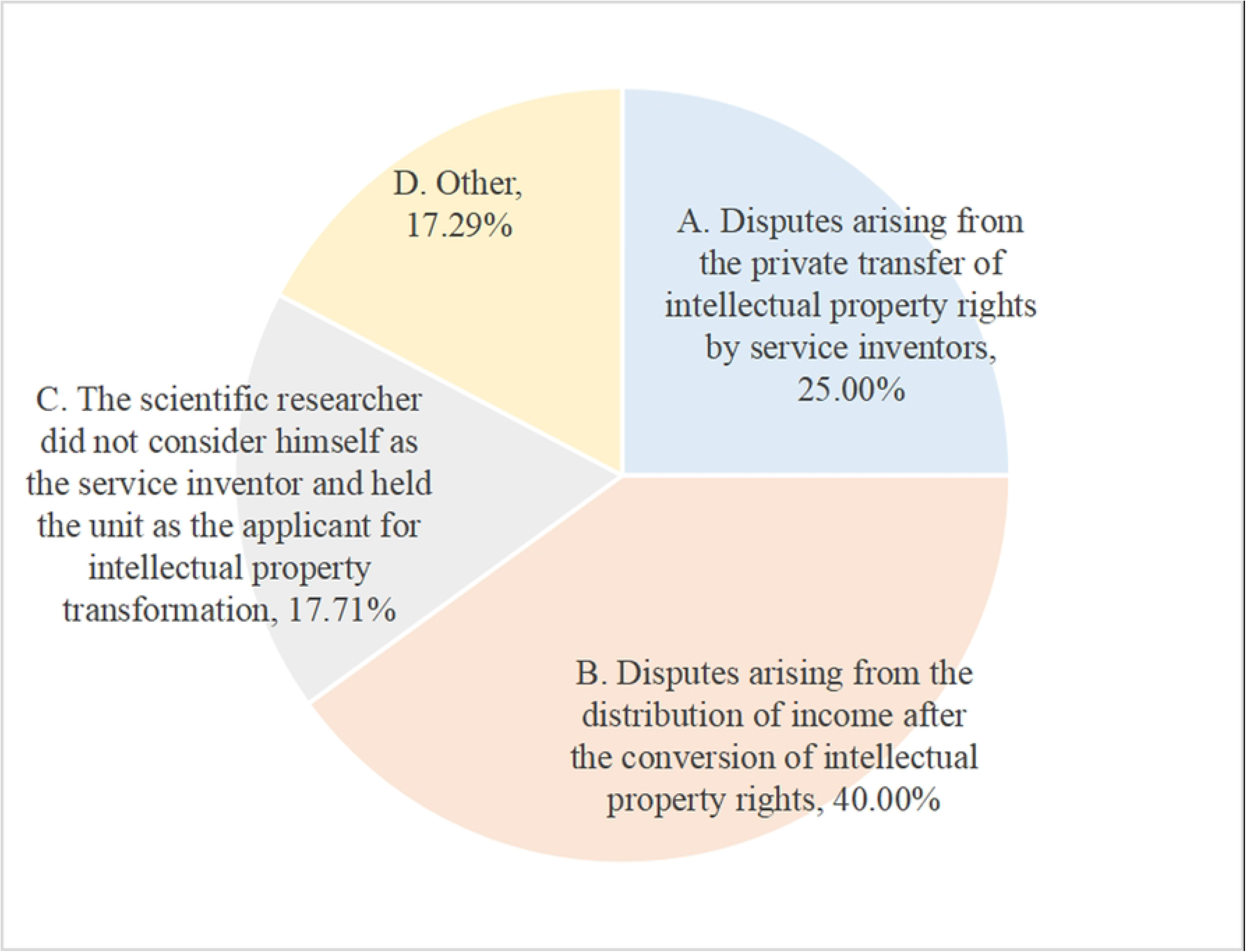
What are the main disputes between your university and scientific researchers during transformation of STA? STA=science and technology achievement

**Fig 19.**
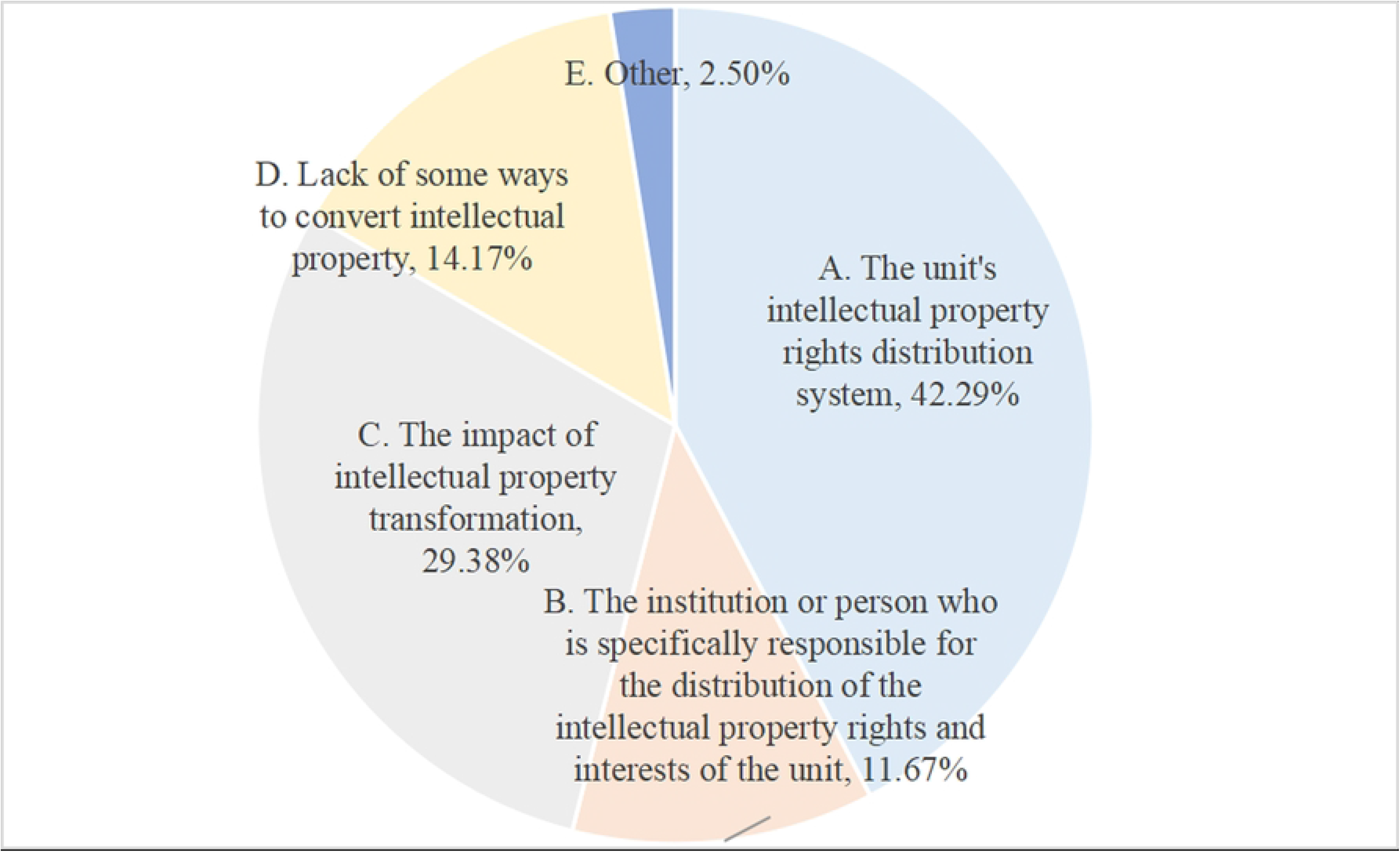
What is the main obstacle to the distribution of RISTA? RISTA=rights and interests in science and technology achievements

**Fig 20.**
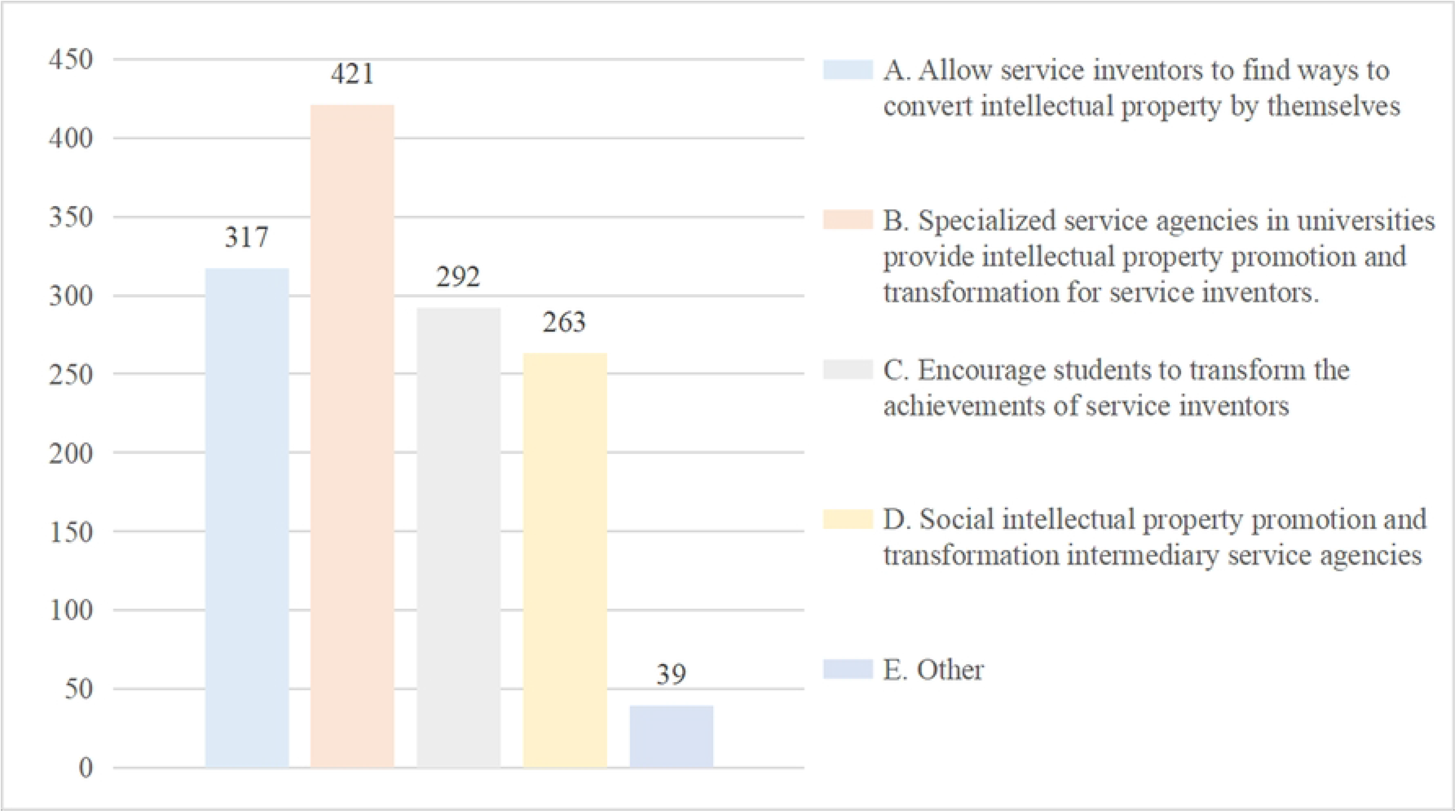
What are the ways to promote the transformation of STA ? STA=science and technology achievement

**Table 1.** Demographic summary of participants.

| **Demographic variables** |  | **Frequency (n)** | **Percentage (%)** |
| --- | --- | --- | --- |
| **Age** | ≤27 | 113 | 33.24 |
|  | 28-31 | 41 | 12.06 |
|  | 32-35 | 41 | 12.06 |
|  | 36-39 | 41 | 12.06 |
|  | 40-50 | 41 | 12.06 |
|  | > 50 | 63 | 18.53 |
| **Education** | junior college | 0 | 0.00 |
|  | undergraduate | 41 | 12.06 |
|  | master | 134 | 39.41 |
|  | doctor | 165 | 48.53 |
|  | other | 0 | 0.00 |
| **professional title** | elementary | 21 | 6.18 |
|  | intermediate | 41 | 12.06 |
|  | vice-senior | 103 | 30.29 |
|  | senior | 52 | 15.29 |
|  | other | 123 | 36.18 |
| **Profession** | Researchers in the field of intellectual property (such as intellectual property, technology transfer, management, and other majors or fields) | 144 | 42.35 |
|  | Intellectual property management department staff/technology transfer agency staff/patent examiners | 52 | 15.29 |
|  | Inventors/technical personnel (such as scientific and technological researchers in colleges and universities, etc.) | 103 | 30.29 |
|  | others | 41 | 12.06 |
| **length of employment** | < 1 | 82 | 24.12 |
|  | 1-3 | 72 | 21.18 |
|  | 4-6 | 31 | 9.12 |
|  | 7-9 | 21 | 6.18 |
|  | ≥10 | 134 | 39.41 |
| Total |  | 340 | 100.00 |

## Result

Combined with the relevant literature [22–26] and the results of the questionnaire survey, it is found that the transformation of scientific and technological achievements in China’s universities mainly exists in the following problems:

### Defects in the management system for the distribution of rights and interests of scientific and technological achievements

In the process of transforming scientific and technological achievements in colleges and universities, the most important dispute between universities and scientific researchers is “disputes caused by the distribution of benefits after the transformation of scientific and technological achievements”, and most university teachers believe that the biggest obstacle hindering the realization of better distribution of intellectual property rights and interests is “the unit’s scientific and technological achievements rights and interests distribution system itself”, and reflects that the incentive effect of the rights and interests of scientific and technological achievements of universities for scientific and technological researchers is poor. It cannot really play a positive role in mobilizing the scientific and technological achievements created by scientific researchers to develop in the direction of the transformation of scientific and technological achievements. The main problems are as follows:

First, the loss of intellectual property rights in some universities is relatively serious, and there is a lack of a reasonable distribution and management system for the rights and interests of scientific and technological achievements. According to the survey, 77% of universities have introduced a management system for the distribution of rights and interests in scientific and technological achievements in advance, but more than 20% of universities have not issued relevant management systems. The management system is lacking, and scientific researchers have no laws and regulations to rely on in terms of the proportion of post scientific and technological achievements, private transfer of intellectual property rights, and intellectual property rights holders, which has led to disputes over the rights and interests of universities and scientific researchers.

Second, the distribution of rights and interests in scientific and technological achievements in most colleges and universities does not take into account of the need of the rights and interests of the persons with scientific and technological achievements. In terms of the attribution of patent results, 98% of the researchers said that the unit did not issue a policy provision that the person of post scientific and technological achievements can become the right holder together with the unit, while 93% of the researchers believe that the common right holder is not only “a kind of rights protection for the person of the post scientific and technological achievements”, but also “helps the person of the post scientific and technological achievements to devote himself to scientific research”, so it is necessary for the person of the post scientific and technological achievements of intellectual property rights to become the right holder together with the unit. In terms of the distribution of benefits, 73% of the researchers expressed their willingness to start their own high-quality intellectual property rights (fig 21), but hoped that the proportion of universities in the equity structure was less than 30% (fig 22), and the survey showed that 75% of the researchers reflected that the proportion of post scientific and technological achievements of university scientific and technological achievements was less than 70%, which was different from the expectation that post scientific and technological achievements required to account for more than 70% of the share capital, hindering the transformation of scientific and technological achievements.

**Fig 21.**
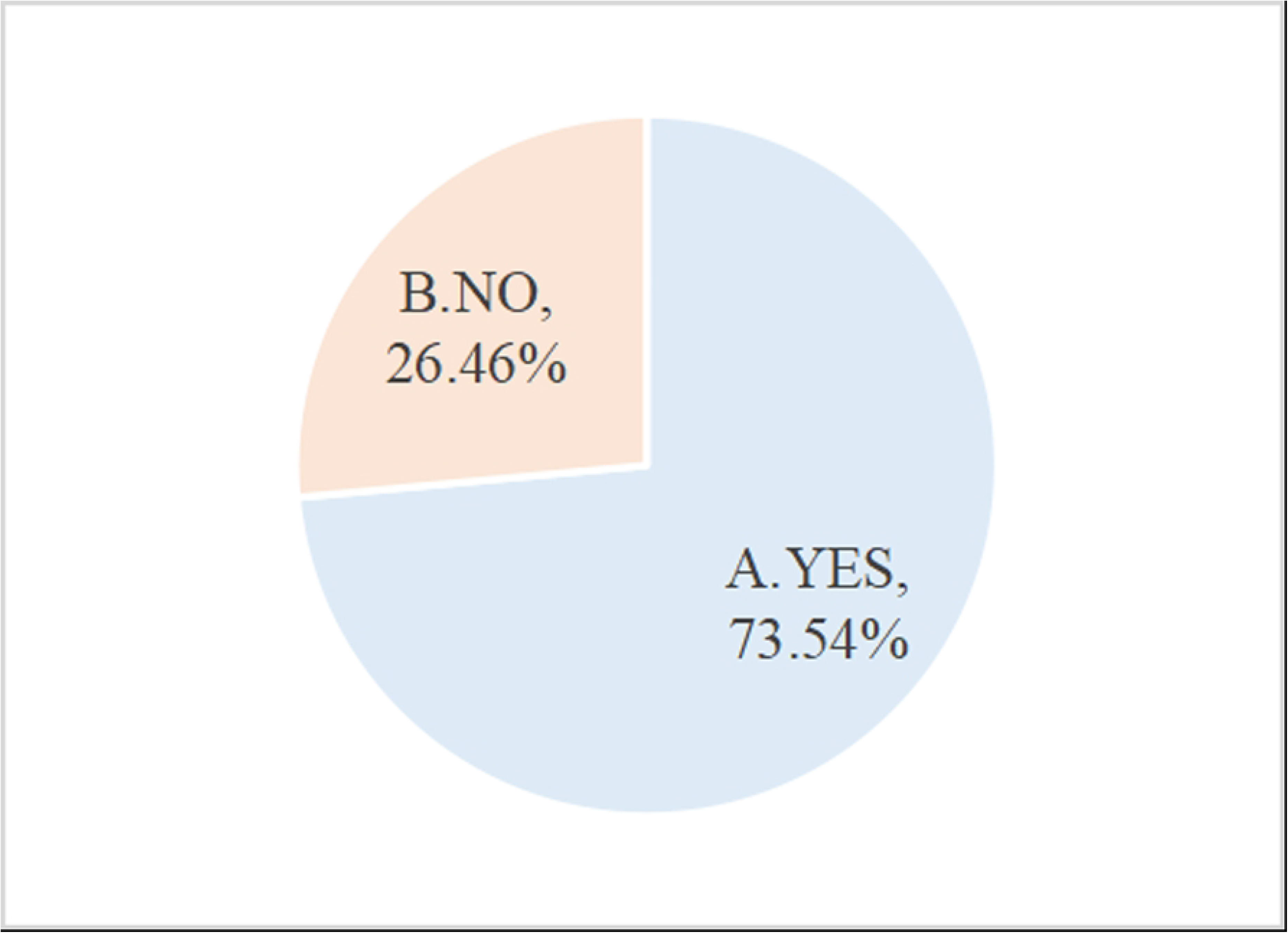
Are you willing to start an undertaking with your high-quality intellectual property?

**Fig 22.**
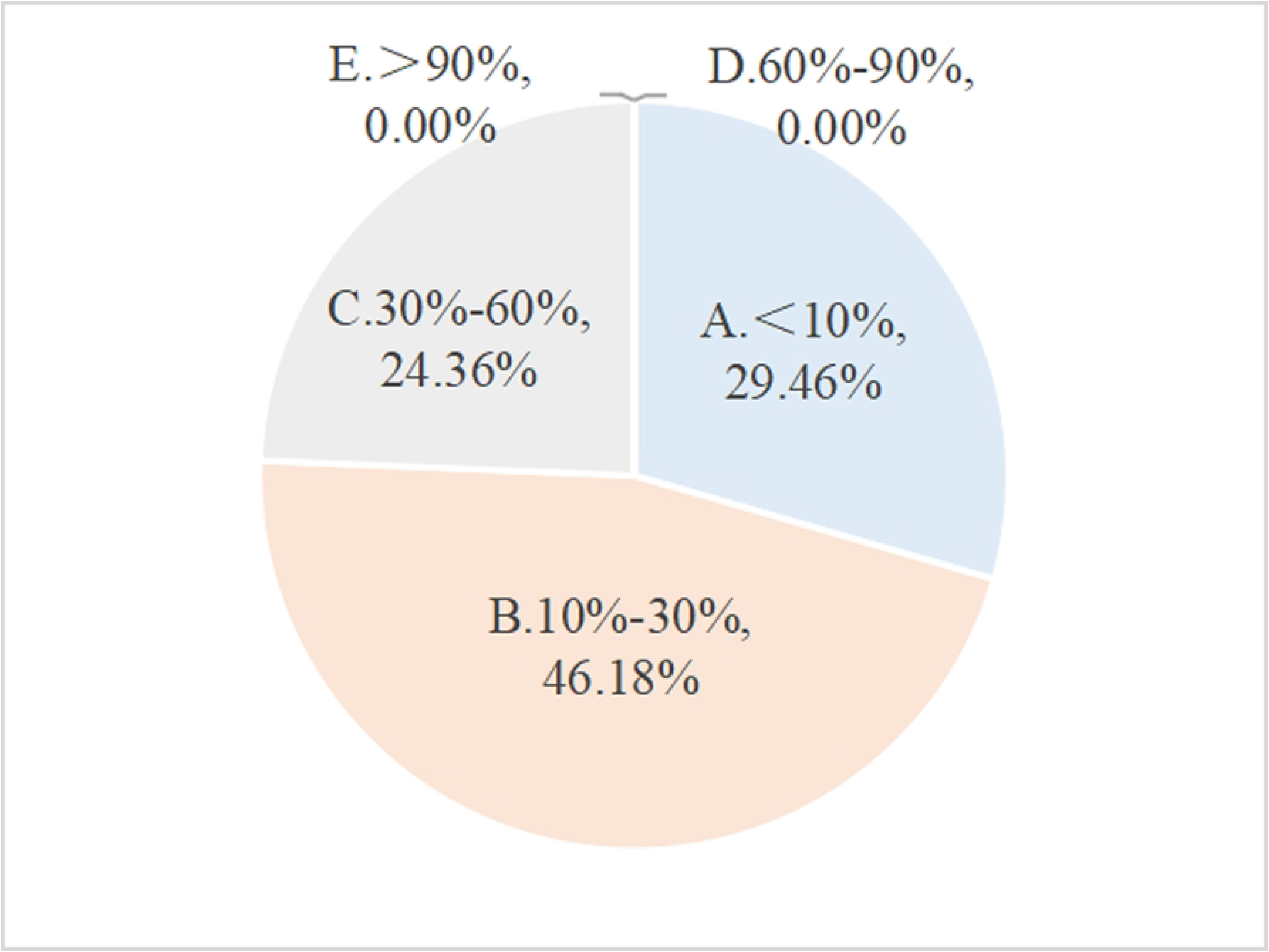
What is the percentage of the university’s share in capital structure that you want?

Third, most universities have many deficiencies in publicizing and implementing the management system for the distribution of rights and interests in scientific and technological achievements. According to the survey, 56% of the researchers said that they did not understand the management system for the distribution of rights and interests in scientific and technological achievements issued by their units, indicating that universities did not fully publicize the management system for the distribution of rights and interests in scientific and technological achievements. In the absence of understanding the distribution management system, scientific researchers are prone to disputes with universities in terms of intellectual property transfer, distribution of income from the transformation of achievements, and intellectual property rights holders. Due to the existence of the distribution management system, the university is in an advantageous position in disputes, and scientific researchers sometimes have to choose to give in.

### Imperfect evaluation indicators of scientific and technological achievements

The project assessment mechanism of colleges and universities mostly adopts the workload recognition scoring system, and the assessment scope includes: decision-making consultation, technical services, technical consultation, etc. The benefits of technological achievement transformation are not included in the evaluation scope, while the main purpose of 43% of researchers applying for patents is to complete the assessment indicators, but because the transformation benefits of scientific and technological achievements are not helpful to complete the assessment indicators, the scientific and technological achievements have not been further transformed. In terms of scientific research awards, in order to better stimulate the enthusiasm and creativity of the majority of scientific researchers, most universities adopt a horizontal scientific research project to the account funding incentive system, but due to the low reward of invention patents and the school no longer take out additional funds as a reward, resulting in a lack of enthusiasm for scientific researchers.

In the evaluation of professional titles, taking the qualification conditions of professors in a province as an example, “based on their own work, giving full play to their own professional advantages, engaged in scientific and technological development, scientific and technological achievements promotion work, earned remarkable achievements, achieved huge economic benefits and gained better social benefits, won the provincial level or above the promotion of scientific research achievements commendation, or obtained more than 2 national invention patent authorization.” There are no specific quantitative indicators for the economic and social benefits achieved by the transformation of the results, making the indicators of the transformation of the results useless. Although the current policy of identifying relevant titles, that is, the evaluation of excellence and evaluation of the first, mentions the transformation of scientific and technological achievements, but in the process of policy implementation, it is far less than the role of papers and other scientific and technological achievements, which has caused universities to produce a large number of low-value scientific and technological achievements. According to the above survey, the existing scientific researchers have a high voice for the quantitative indicators of the transformation of the achievements into the assessment indicators, and 79% of the researchers expressed their willingness to link the transformation effect of the scientific and technological achievements of the post with the assessment indicators (projects, titles, evaluations, etc.). Therefore, when implementing the mixed ownership system of scientific and technological achievements in actual posts, colleges and universities also need to adjust and improve the assessment and evaluation policies for scientific and technological achievements from the university level.

### Imbalance in the service system for the transformation of scientific and technological achievements

70% of the researchers reflect that the unit has established a technology transfer agency or a third-party institution to carry out technology transfer services, 65% reflect that there is a special intellectual property agency or a third-party agency as an intellectual property agency, while only 44% reflect that there is a special scientific and technological achievement value assessment agency or a third-party institution for value assessment, and 52% reflect that there are special scientific and technological achievement transformation service institutions or third-party institutions for scientific and technological achievements transformation services. Universities pay much attention to technology transfer and intellectual property agency services, but there is a lack of value assessment and achievement transformation services, and there is an imbalance in the construction of the service system for the transformation of scientific and technological achievements, so it is necessary to appropriately enhance the work in value assessment and achievement transformation services to balance the intellectual property service system.

### 2.4 Lack of authoritative standards for the evaluation of the value of the transformation of scientific and technological achievements

When the transformation of scientific and technological achievements is carried out by universities, the most important way to determine the transformation value is “the transfer party agrees to discuss the value”, which lacks professional, scientific and rational evaluation methods due to the use of discussion to determine the value of the transformation of scientific and technological achievements, and in most cases, there is a difference of more than twice the expected value of intellectual property rights and the value of actual transformation by researchers. The main reason for the large difference between the expected value and the actual conversion value is the lack of “legal and regulatory support from the government or school” and “the support of the university’s own achievement transformation institutions (or entrusted third-party intermediary service agencies)”.

## Solutions

### Formulate a mixed ownership management system for scientific and technological achievements in colleges and universities

Each university shall formulate the Specification for the Mixed Ownership Management System of Scientific and Technological Achievements of Colleges and Universities according to its own actual situation.

Implement the standardization of the equity distribution system of mixed ownership of post scientific and technological achievements. As shown in Fig 23, the implementation path of the mixed ownership equity distribution model of post scientific and technological achievements is constructed by comprehensively considering the factors of service invention results in the distribution of rights and interests[31], such as the contribution of each subject in the generation and transformation of service invention achievements, and to build a job scientific and technological achievement transformation model led by employers, scientific and technological workers and service agencies, so as to promote the effective agglomeration of resources under the innovation ecosystem and the synergy between various subjects. Through game analysis of the process, proportion and transformation decision of the multi-stakeholder parties of the mixed ownership of the service invention results, a feasible optimization equity distribution path is found, and a dynamic equity distribution model standard is established. The selection of the service invention achievement model under the leading mode of different subjects is analyzed, and finally the implementation system of the dynamic distribution model of mixed ownership rights and interests of service inventions is formed relative to the different subject leading models. At the same time, in order to reflect the pertinence of the system more, colleges and universities can also consult scientific researchers on the distribution intentions of post scientific and technological achievements such as the attribution and transformation of scientific and technological achievements through employee congresses, research questionnaires, etc., determine the proportion of ownership of post scientific and technological achievements, the proportion of distribution of transformation benefits, and other issues and incorporate them into the policy documents of universities for implementation.

**Fig 23.**
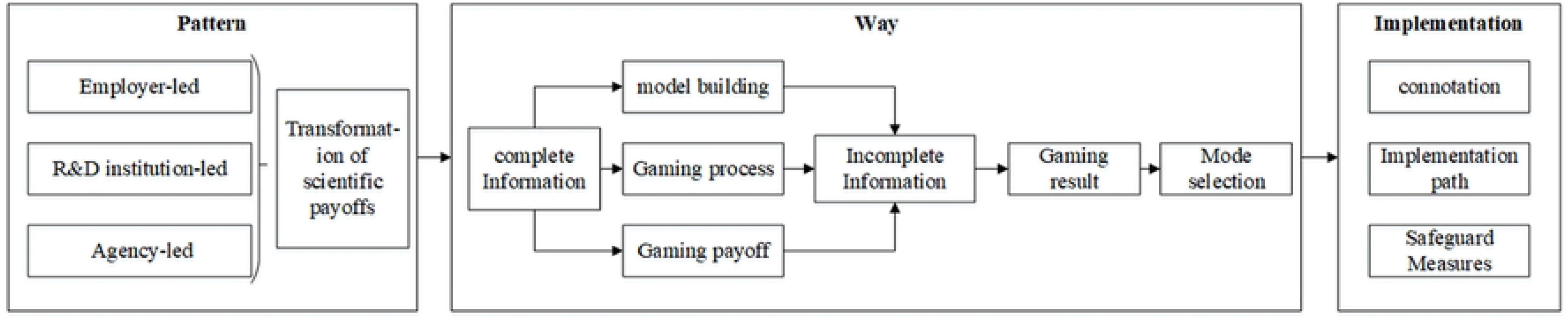
Implementation path of the mixed ownership rights distribution model for job science and technology achievements.

Explore the construction of a mixed ownership culture of scientific and technological achievements in colleges and universities. Through tangible institutional arrangements, policy guidance and reasonable incentive measures, the construction of mixed ownership culture of scientific and technological achievements in colleges and universities is carried out to improve the enthusiasm of those who have completed scientific and technological achievements to apply for patents and participate in the subsequent implementation of patents. Colleges and universities can actively carry out the transformation of scientific and technological achievements by building a mixed ownership cultural wall of post scientific and technological achievements, regularly update them, and include the transformation of scientific and technological achievements into the scope of performance appraisal and professional title evaluation and assessment to stimulate university researchers to actively participate in the transformation of scientific and technological achievements.

Strengthen the management of the mixed ownership of scientific and technological achievements in colleges and universities. When determining the proportion of ownership distribution and the proportion of transformation income distribution of post scientific and technological achievements, university administrators should strictly implement it in accordance with the relevant systems for the distribution of ownership and the distribution of transformation benefits. In terms of transformation of scientific and technological achievements in colleges and universities, the scientific research management department must carefully analyze and review the terms of the contract in the early stage, strictly control and supervise the implementation of the contract, and conduct stage evaluation and experience summary of the effectiveness of the later stage of transformation. Colleges and universities should set up full-time posts, introduce professional talents or entrust professional third parties in the evaluation of scientific and technological achievements and the transformation of scientific and technological achievements, carry out professional docking, and strengthen process management.

### Improve the mixed ownership assessment system for scientific and technological achievements in colleges and universities

All colleges and universities should improve the Specification of the Mixed Ownership Assessment System for Scientific and Technological Achievements of Colleges and Universities in accordance with national and local laws and regulations.

In the process of internal reward assessment of colleges and universities, the traditional system of taking the number of papers and patents as the standard should be broken, and a multi-dimensional assessment method should be formulated based on academic and theoretical level, the quality of papers and the transformation of patent technology and economic benefits. The reward mechanism of colleges and universities is based on scientific research topics, papers, patent applications, authorization and as the core of the promotion and reward mechanism [28], taking a university in Jiangsu Province as an example, the scientific research topics, papers, patent applications and authorizations are exchanged for corresponding scores, and each year the school assesses the colleges, requiring them to achieve a certain score, and the colleges will send the total score task of the colleges to the annual assessment indicators of researchers at all levels, forcing scientific researchers to be oriented to the academic nature of scientific and technological achievements, rather than application. As a result, the school’s scientific researchers have gradually attached importance to the publication of papers and patent applications, resulting in a lot of low-quality scientific and technological achievements, and also seriously affecting the enthusiasm of the school’s scientific and technological personnel to engage in the practical application and promotion of scientific and technological achievements. Therefore, it is necessary to change the existing incentive policy of “heavy topic papers, heavy patent authorization, and light patent implementation”.

According to the characteristics of the mixed ownership of post scientific and technological achievements, improve the existing professional title promotion methods, awards and appraisals, post appointment implementation measures and other relevant management documents of colleges and universities, and select the best policies to absorb and mobilize the majority of scientific research personnel to participate in the transformation of science and technology. In the evaluation of the titles of scientific research personnel, the amount of mixed ownership of the rights and interests of the scientific and technological achievements of the post should be comprehensively considered, and the benefits of the actual transformation of scientific and technological achievements should be paid attention to, and even greater than the weight given by scientific research topics and papers. In terms of awards and appraisals, the school can take out part of the benefits of the transformation of scientific and technological achievements, and set up two types of scientific and technological awards to reward scientific researchers and outstanding scientific and technological service personnel who pay attention to the transformation of scientific and technological achievements. In terms of post appointment, conduct full-time guidance for employees engaged in scientific and technological services, and focus on assessing their professionalism and service capabilities in terms of post appointment. At the same time, in colleges and universities, we should also pay attention to the scientific research and development personnel engaged in the research and development of basic and cutting-edge advanced technologies, the basic talents engaged in the research and development of cutting-edge advanced technologies and the applied talents who pay attention to the transformation of scientific and technological achievements, formulate different categories of classification assessment indicators for research and development personnel, fully mobilize the innovation awareness and enthusiasm of various types of scientific research personnel in schools, and realize the value of scientific research personnel and the value of transfer and transformation.

### Implement a mixed ownership service system for scientific and technological achievements in colleges and universities

All colleges and universities should actively build relevant organizational structures and implement the Specification of Mixed Ownership Service System for Scientific and Technological Achievements of Colleges and Universities.

Construct a high-quality and efficient ownership service office for scientific and technological achievements. The transformation of scientific and technological achievements in colleges and universities is a complex process of technological and economic change from the production of scientific and technological achievements to the commercialization of scientific and technological achievements [27]. The transformation of scientific and technological achievements of colleges and universities must start from rigorous laboratory research records, followed by pre-management links such as disclosure and quality assessment of scientific and technological achievements, and finally enter the implementation and management of the transformation of scientific and technological achievements. Therefore, a mixed ownership service system for scientific and technological achievements in colleges and universities should be built from the whole process of the generation and transformation of scientific and technological achievements. The service system of the specific scientific and technological achievements ownership service office includes: First, build an information bridge through the office and disclose the scientific and technological achievements of scientific researchers; then, the office assists universities and teachers to carry out the quality and value assessment of the scientific and technological achievements of the job, and the evaluation work can be carried out by the office or the high-quality third-party service agency that cooperates; Secondly, establish an extensive cooperation network through the office in the strategic integration of scientific and technological achievements of universities and scientific and technological achievements after the professional evaluation of potential buyers, and assist researchers to undertake relevant negotiations and process procedures in the transaction of scientific and technological achievements; Finally, it is necessary to formulate relevant assessment measures for inventors of scientific and technological achievements to participate in the specific industrialization process of scientific and technological achievements, and office staff should also assist inventors of scientific and technological achievements to negotiate relevant contract matters such as the distribution of relevant benefits with the transformed enterprises, promote the transformation of scientific and technological achievements, and reduce the risk of failure of achievement transformation.

Build a scientific value assessment expert think tank team. Universities can carry out expert classification according to the industry, build a patent value evaluation expert think tank, and use the advantages of technical experts on campus to carry out authoritative assessment of patent quality and value. Strengthen the screening and judgment of high-quality and high-value patents, and the service offices or high-quality service institutions of scientific and technological achievements of universities should initially screen out funny high-quality and high-value patents from the perspective of the technology market and the application market. To formulate flexible valuation standards for scientific and technological achievements, the ownership service office of scientific and technological achievements of universities or high-quality service institutions need to formulate appropriate value evaluation methods through traditional evaluation methods such as market method, revenue method, cost method, industrial law, etc., combined with diversified methods such as fuzzy neural network method of real option method, and according to the perspective of the core key technology of the patent, market prospects, and legal authority.

Actively build a diversified accelerator and incubator platform to promote the landing of job scientific and technological achievements. As an important population of innovative development communities in the innovation ecosystem, various accelerators and incubators are an important boost to further develop and commercialize the scientific and technological achievements of universities and establish start-ups or derivative enterprises. Colleges and universities should establish a certain number of accelerators and incubator platforms to meet the needs of teachers and students to establish start-ups or spin-offs through scientific and technological achievements. Meanwhile, it can integrate the start-ups or spin-off companies that are cultivating or have been formed, establish a university science and technology park, drive the scientific and technological and economic development of the area near the university, and promote the formation of a regional innovation ecosystem.

## Conclusion

Compared with other state-owned enterprises and institutions, the scientific and technological achievements of private enterprises and universities are concentrated in basic research, and rarely consider market demand, and there is no requirement for compulsory commercialization, so most of them are concluded in the form of papers. College teachers and scientific research teams apply for projects with the utilitarian purpose of promoting titles and reporting awards. It is recommended to start from the division of property rights and rights and interests of scientific and technological achievements, by encouraging the completion of job scientific and technological achievements to participate in the whole process from project establishment to the transformation of scientific and technological achievements, to achieve the full use of job scientific and technological achievements, improve the innovation enthusiasm of inventors to drive scientific and technological development, improve the supply-side reform of scientific and technological achievements, reduce the loss of valueless state-owned assets, and combine the reward problem of completers.

The original intention of the mixed ownership of scientific and technological achievements in colleges and universities is to solve the problem of ownership and distribution of benefits of scientific and technological achievements, with a view to distributing the benefits of scientific and technological achievements among universities and job scientific and technological achievements, and forming a patent common between universities and job scientific and technological achievements. But there are practical and theoretical obstacles. The obstacles encountered at the institutional level are mainly defects in the management system for the distribution of rights and interests of scientific and technological achievements, that is, the Patent Law only stipulates the ownership of the units of the job scientific and technological achievements, and does not stipulate the ownership of the rights of the person who completes the job scientific and technological achievements, and only makes the minimum value for the distribution of the benefits of the transformation (the extraction of the conversion royalties is not less than 10%), and the minimum value is much less than the value created by the person who completes the job scientific and technological achievements. At the operational level, it is mainly due to the relatively backward management thinking, resulting in imperfect evaluation and assessment indicators of scientific and technological achievements, imbalance in the service system for the transformation of scientific and technological achievements, and lack of authoritative standards for the evaluation of the value of the transformation of scientific and technological achievements. Therefore, each university should formulate the Specification for the Mixed Ownership Management System of Scientific and Technological Achievements of Colleges and Universities according to its actual situation, implement the standardization of the rights and interests distribution system of mixed ownership of job scientific and technological achievements, explore the construction of mixed ownership culture of scientific and technological achievements of colleges and universities, and strengthen the management of the mixed ownership of scientific and technological achievements of colleges and universities. All colleges and universities should improve the Specification of the Mixed Ownership Assessment System for Scientific and Technological Achievements of Colleges and Universities in accordance with national and local laws and regulations, and in the process of internal reward assessment of colleges and universities, they should break the traditional system of taking the number of papers and patents as the standard, and formulate multi-dimensional assessment methods based on academic and theoretical levels, the quality of papers and the transformation of patented technology and economic benefits. According to the characteristics of the mixed ownership of job scientific and technological achievements, improve the existing professional title promotion methods, appraisal and appraisal methods, post appointment implementation measures and other relevant management documents of colleges and universities, and select the best policies to absorb and mobilize the majority of scientific research personnel to participate in scientific and technological transformation work. All colleges and universities should actively build relevant organizational structures, implement the Specification for the Mixed Ownership Service System for Scientific and Technological Achievements of Colleges and Universities, build a high-quality and efficient ownership service office for scientific and technological achievements, build a scientific value assessment expert think tank team, and actively build a diversified accelerator and incubator platform to promote the landing of job scientific and technological achievements. The implementation of the mixed ownership implementation system for scientific and technological achievements of posts in colleges and universities is conducive to the implementation of the reform of mixed ownership of job science and technology achievements, and is also conducive to the effective transfer and industrialization of post scientific and technological achievements.

It is believed that with the continuous implementation of the mixed ownership reform of scientific and technological achievements of colleges and universities, more problems will be solved. Therefore, in the future work and study, the author will conduct in-depth research on new problems arising from the practice of mixed ownership of scientific and technological achievements, put forward corresponding countermeasures and suggestions, and continuously improve the relevant content of mixed ownership of scientific and technological achievements in colleges and universities.

## Author Contributions

**Conceptualization:** Panjun Gao, Yong Qi

**Data curation:** Panjun Gao, Qing Guo

**Formal analysis:** Panjun Gao

**Funding acquisition:** Yong Qi

**Investigation:** Panjun Gao, Qing Guo

**Methodology:** Panjun Gao

**Project administration:** Yong Qi

**Resources:** Yong Qi

**Software:** Panjun Gao

**Supervision:**Yong Qi

**Validation:** Panjun Gao, Qing Guo

**Visualization:** Panjun Gao

**Writing – original draft:** Panjun Gao

**Writing – review & editing:** Yong Qi, Qing Guo

